# Translational Activity Controls Ribophagic Flux and Turnover of Distinct Ribosome Pools

**DOI:** 10.1101/2022.05.13.491786

**Authors:** Jakob Trendel, Milan Aleksić, Matilde Bertolini, Marco Jochem, Günter Kramer, Stefan Pfeffer, Bernd Bukau, Jeroen Krijgsveld

**Author notes:** Correspondence to: Jeroen Krijgsveld, German Cancer Research Center (DKFZ), Im Neuenheimer Feld 581, 69120 Heidelberg, Germany.

## Abstract

Ribosomes are among the most abundant and complex machineries in the cell, however, the turnover of their subunits remains poorly understood. Here, we apply proteomic flux and cryo-electron microscopy analyses to interrogate the ribosome life cycle in human cells. We show that subpopulations of ribosomal subunits coexist, which vary in turnover kinetics and structure. Specifically, 80S ribosomes have a much longer half-life than free 40S and 60S ribosomal subunits, indicating that they represent distinct subunit pools that poorly intermix. Translation inhibition starkly increases the pool-size of 80S ribosomes in a translationally idle state and induces ribophagy of old ribosomes, ultimately rejuvenating the ribosome fleet. Our findings provide a comprehensive model for ribosome turnover and its regulation via translational activity.

## Introduction

Cells continuously turn over their proteome through protein biosynthesis and decay, where ribosomes constitute the core machinery translating messenger RNA (mRNA) into nascent proteins, while the ubiquitin-proteasome and the autophagy-lysosome system are the main pathways for protein degradation ^1^. Ribosomes are highly abundant, representing six percent of the protein mass in mammalian cells, and more than 80 percent of the total RNA ^2^. Throughout their lifetime eukaryotic ribosomal proteins can be part of multiple assembly states. A large excess of free ribosomal protein is continuously produced in the cytosol and shuttled into the nucleus, only a fraction of which is incorporated into nascent ribosomal subunits, whereas the majority is quickly degraded by the proteasome ^3–6^. Fully assembled small (40S) and large (60S) ribosomal subunits undergo final maturation in the cytosol ^7^ before they combine into a translation-competent 80S complex at the start codon of mRNA during translation initiation ^8^. To release the ribosome from mRNA upon arrival at the stop codon, the 80S complex must be split into small and large subunits which then join the pool of free subunits ^9^, although immediate re-joining of subunits can occur on the same transcripts in another round of initiation in a process called ribosome recycling ^10,11^. Additionally, a significant proportion of subunits is kept in translationally inactive 80S complexes (also known as idle, empty or hibernating ribosomes), whose origins, fate and function remain elusive. The proportion of idle 80S ribosomes strongly increases upon serum withdrawal, which has been utilized for the preparation of pure 80S populations for structural studies ^12,13^. Degradation of ribosomal complexes, especially under nutrient-poor conditions, is mediated by ribophagy, a selective form of autophagy ^14–17^. Ribophagy receptors such as the human NUFIP1 selectively recruit ribosomes to autophagosomes ^18^ replenishing the cellular nucleotide and amino acids pools in times of starvation ^19^. In addition, we recently reported a rapid form of ribophagy during stress-induced translational arrest, which eliminates up to 50 % of ribosomal protein within 30 minutes ^20^. This fast form of ribophagy only minutes after stress is markedly different to long term degradation events that happen over the course of several hours to days during nutrient withdrawal.^15,17^ Despite detailed insight in ribosome biogenesis and function ^7–9^, fundamental questions remain towards how ribosome homeostasis is maintained, and how turnover of a ribosomal protein differs before and after becoming part of a ribosomal complex. In addition, it is unknown if ribosome biogenesis and ribophagy are independent or functionally coupled processes, and if translational activity influences these turnover kinetics. To address these questions and determine how protein flux is regulated in ribosomal complexes, we applied two orthogonal pulsed-SILAC proteomic approaches that revealed the lifecycle of ribosomal subunits along different assembly states in human MCF7 cells. Using cryo-electron microscopy (EM), we show that virtually all ribosomes outside the polysome fraction are translationally inactive. Furthermore, we demonstrate that different forms of translational arrest strongly increase the pool of idle 80S ribosomes, triggering ribophagy. Collectively, our findings demonstrate that ribosome homeostasis is intimately linked to translational activity.

## Results

### A Highly Robust Normalization Procedure for Protein Half-Life Measurements within Purified Ribosomal Complexes

In turnover measurements of the total proteome all possible assembly states of a ribosomal protein are averaged into one protein half-life, so that information about the turnover within each state is lost. In order to compare the half-lives of ribosomal proteins in different ribosomal complexes we purified them individually from MCF7 cells with polysome profiling. Therefore, we first verified that this could be performed reproducibly and quantified proteins within the individual fractions using intensity-based, label-free proteomics (Fig. S1A). Indeed, we found good precision of our MS analysis between biological replicates (Fig. S1B, Table S1) and reproduced the stoichiometric range of ribosomal proteins anticipated from earlier work ^21^. To derive protein half-lives within ribosomal complexes, we combined polysome profiling with a time course of pulsed-stable-isotope-labelling in cell culture (pulsed-SILAC) multiplexed with tandem-mass-tag labelling (TMT) (Fig. 1A). This labelling strategy was initially presented as iTRAQ-4Plex-SILAC ^22^, first implemented in human cells as TMT-10Plex-SILAC ^23^ and recently revisited by Zecha et al., who greatly improved the analysis of TMT-SILAC data using a single search in MaxQuant ^24^. To minimize experimental variability, we adapted the TMT-SILAC approach by Zecha et al. by changing the SILAC labelling scheme to allow for the processing of identical amounts of cells from all time points in one common ultracentrifugation run for polysome profiling. Therefore, SILAC heavy-labelled MCF7 cells were switched to SILAC light media at nine time points between 32 to 2 hours before the simultaneous cell harvest (Fig. 1A). Indeed, this resulted in a reproducible fractionation of ribosomal complexes for all time points, albeit with remaining variability of the amounts contained within each fraction (Fig. 1B). Subsequent MS quantification of their protein constituents showed strong discontinuity between time points, impeding meaningful derivation of ribosomal protein half-lives from their decay curves (illustrated for 80S ribosomes in Fig. 1C, left). Using a similar experimental setup to derive protein half-lives for the total proteome of MCF7 cells we observed much better data continuity (Fig. S1D), indicating that the extensive sample processing required for the purification of ribosomal complexes had introduced strong variation to our MS measurements that could not be avoided despite our efforts to process equal amounts of all samples in one batch. Conceptually, the decay model applied here to derive protein half-lives requires that protein amounts stay constant across time points. Previous studies approached this by normalizing the combined TMT reporter intensities of each time point to one common total sum that is identical for all TMT channels (total sum normalization, TSN) ^23,24^, introducing the assumption that the total amount of protein is constant across time points if the combined TMT reporter intensity is constant across TMT channels. Applying this normalization to our data (Fig. 1C left, Fig. S1D, H) or published data from Zecha et al. (Fig. S2A) we noticed that for individual peptides the resulting sum of TMT reporter intensities between the light and heavy SILAC channel was often very inconsistent between time points. For peptides of the same protein this inconsistency prevailed even within one time point and, additionally, differed between replicates (Fig. 1D, top).

**Fig. 1:**
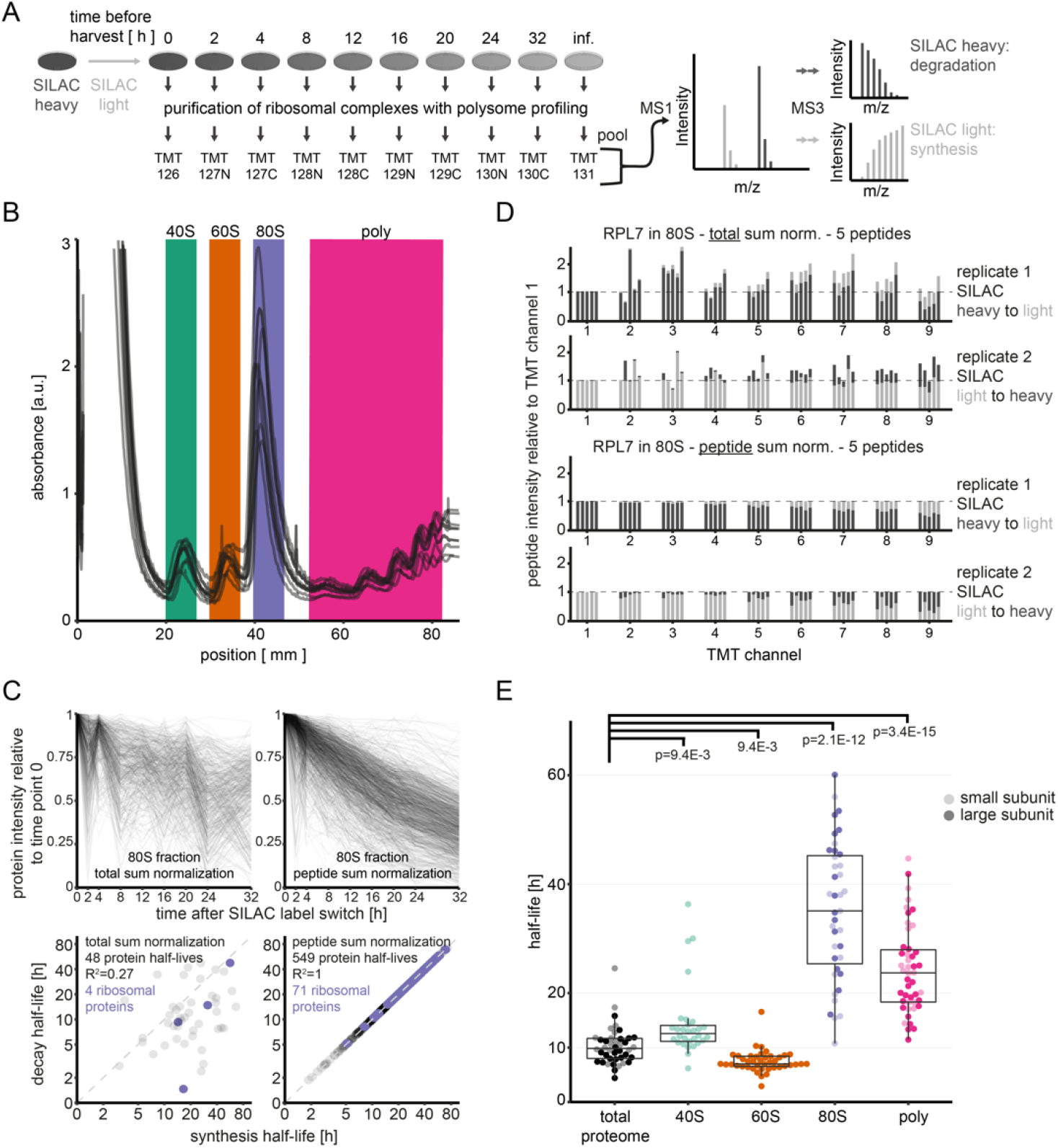
Comparing protein half-lives between the total proteome and ribosomal complexes. A) MS3-based pulsed-SILAC-TMT strategy for the determination of protein half-lives. SILAC heavy-labelled cells are switched to SILAC light media to monitor protein degradation (SILAC heavy) and synthesis (SILAC light) within ribosomal complexes over time. B) Polysome profiling demonstrating reproducibility for the purification of ribosomal assemblies from MCF7 cells. Traces represent the nine time points during a pulsed-SILAC experiment, and colors indicate fractions that were collected to perform MS analysis and derive protein half-lives. C) Comparison of total sum (left panels) and peptide sum normalization (right panels) for the determination of protein half-lives from purified ribosomal complexes. Displayed is exemplary data for the 80S fraction as purified in B from MCF7 cells. Top panels show protein decay curves used for model fitting and computation of decay half-lives below. Bottom panels show scatterplots comparing protein synthesis and decay half-lives with purple coloring indicating ribosomal proteins. Note logarithmic scaling. D) Barplots showing the TMT peptide intensities from the light (light grey) and heavy (dark grey) SILAC channels for the five peptides of RPL7 quantified in the 80S fraction. Intensities are normalized to TMT channel 1 for comparison. Top two panels show data from two biological replicates with SILAC label-swap normalized with total sum normalization, bottom two panels the same data normalized with peptide sum normalization. E) Boxplots illustrating half-lives of ribosomal proteins across different cellular contexts. Displayed are means between biological duplicates with SILAC label swap. Proteins known to be part of the small ribosomal subunit are colored pale, large subunit proteins bold. For the 40S fraction only small subunit proteins are shown, for the 60S fraction only large subunit proteins (see also Fig. S1B). Pairwise testing occurred with a Wilcox ranksum test for proteins shared between the total proteome and each of the other experiments as displayed.

In order to improve the continuity between time points and comply with the prerequisite of the decay model that combined protein amounts must stay constant over time, we introduced peptide sum normalization (PSN), which, for each peptide, normalizes the combined TMT reporter intensities from both SILAC channels to be equal across time points (Fig. 1D, bottom). Applying PSN to TMT-SILAC data of the MCF7 total proteome demonstrated that it was highly beneficial for data continuity, leading to a much larger number of curves that could be used for model fitting with much better fits (Fig. S1E). Consequently, the number of half-lives derived for both protein synthesis and decay in the total proteome of MCF7 cells more than doubled from 921 under TSN (Fig. S1D) to 2402 under PSN (Fig. S1E, Table S2). We validated PSN on published protein decay data from HeLa cells ^24^ where we were able to derive decay as well as synthesis half-lives for 4932 proteins, compared to 3882 proteins using TSN (Fig. S1G, Fig. S2A-E, see Materials and Methods). Moreover, synthesis and decay half-lives were virtually identical under PSN (Pearson R^2^=1), both in our own (Fig S1E) and published data (Fig. S1G) indicating that our new normalization had effectively implemented the necessary prerequisite of the protein decay model that required protein amounts to stay constant over time. This is not a trivial result, since during a TMT-SILAC run protein synthesis is measured independently from protein decay when the mass spectrometer picks a SILAC heavy or light precursor ion to quantify the entire timeline on the MS3 level with TMT reporter ions (Fig. 1A). Interestingly, HeLa protein synthesis half-lives from PSN-corrected data were similar to protein synthesis half-lives from TSN corrected data (R^2^=0.86), and so were protein decay half-lives (R^2^=0.81), whereas within TSN protein decay and synthesis half-lives were rather dissimilar (R^2^=0.42) (Fig. S1G). We tested if PSN could also be applied to different biological replicates and found that cross-replicate normalization (Fig. S1F) performed even slightly better than within-replicate normalization (Fig. S1E), returning 7 % more protein half-lives for the MCF7 total proteome. In addition, we benchmarked PSN by swapping correction factors across time points, consistently obtaining inferior correlation between synthesis and decay, indicating the specificity of PSN (Fig. S2F).

Having established PSN, we applied it to the TMT-SILAC data of the polysome profiling fractions, allowing us to derive hundreds of protein half-lives (1454 from 40S fraction, 1282 from 60S fraction, 549 from 80S fraction, 685 from polysome fraction, Table S2), including most ribosomal proteins (50 from 40S fraction, 79 from 60S fraction, 71 from 80S fraction, 66 from polysome fraction). The decisive improvement effectuated by PSN now made protein half-life measurements in ribosomes possible, as demonstrated for the 80S fraction in Fig. 1C and Fig. S1H&I. In summary, we introduce PSN as a new normalization regimen that significantly improved our own and published half-life measurements of the total proteome, and allowed us to determine protein half-lives within purified ribosomal complexes.

### Protein Half-Lives in Polysome Profiling Fractions Reveal Distinct Pools of Ribosomal Subunits

Using PSN we could now compare protein half-lives between the different ribosomal complexes we had purified using polysome profiling (Fig. 1B), as well as protein half-lives in the total proteome of MCF7 cells. Fig. 1E shows that half-lives of ribosomal proteins were similar in the total proteome and in free 40S and 60S subunits, yet, were strongly stabilized when assembled in 80S complexes of the 80S and polysome fractions. On average ribosomal proteins of the small subunit had 3-fold longer half-lives within the 80S fraction compared to the 40S fraction (p=5.2E-8, Wilcoxon ranksum test), whereas large subunit proteins had 4.6-fold longer half-lives within the 60S fraction compared to the 80S fraction (p=1.0E-10). For the polysome fraction the stabilization was slightly weaker, where small and large subunit proteins were stabilized 1.9-fold (p=1.2E-5) and 3.1-fold (p=1.6E-14), respectively. Notably, small subunit proteins of the 40S fraction were significantly more stable than large subunit proteins of the 60S fraction, explaining the different degrees of their stabilization towards 80S complexes. These observations have three important implications. First, they suggest that free 40S and 60S subunits on the one hand, and 80S ribosomes and polysomes on the other hand, exist as separate pools that poorly mix: If free exchange of subunits existed between the 40S/60S and 80S/polysome fractions, there would eventually be no difference in the half-lives of the ribosomal proteins they contain. Second, since free ribosomal subunits were overall much younger than 80S assemblies, and since ribosome biogenesis is known to produce free 40S and 60S subunits and not 80S assemblies ^7^, this indicates successive aging states along the ribosome life cycle: free 40S and 60S subunits were the youngest, followed by polysomes and finally ribosomes in the 80S fraction. Thus, free small and large subunits have two ways of disappearing from their pool, i.e. being degraded or becoming an 80S complex in the polysome or 80S fraction. Third, by comparing our label-free quantification between MCF7 polysome profiling fractions we estimated that free 40S subunits were twice as abundant as 60S subunits (Fig. S1C, Table S1). The lower abundance and more rapid turnover of 60S subunits suggested that they represented the limiting factor in the formation of new 80S ribosomes (Fig. 1E).

In summary, the half-lives of ribosomal proteins within purified ribosomal complexes revealed different aging stages in the lifetime of ribosomal subunits, and pointed to 80S complexes as a distinct pool characterized by strongly increased stability.

### The Monosome Fraction Predominantly Contains Inactive 80S Ribosomes with a Destabilized P-Stalk Base

Since ribosomes in the 80S fraction exhibited particular stability we investigated if this could be associated with their translational activity or specific structural characteristics. We compared polysome profiles of MCF7 cells at high and low salt, and observed that the vast majority of ribosomes in the 80S peak dissociated into 40S and 60S subunits (Fig. 2A), indicating that they were not attached to mRNA ^25,26^ and were thus translationally idle. To corroborate this, we characterized ribosomes in the pooled non-polysomal fractions by cryo-electron microscopy (cryo-EM) single particle analysis (Fig. S3&4), obtaining a cryo-EM structure of the 80S ribosome at 3.28 Å resolution (Fig. 2B). Analysis of the cryo-EM density confirmed that the 80S ribosomes were translationally inactive, as demonstrated by the complete absence of cryo-EM density for molecular markers of active translation, i.e. mRNA in the ribosomal mRNA channel, a nascent chain in the ribosomal peptide exit tunnel, tRNAs in the ribosomal A-, P- and E-sites (Fig. 2B) or translation elongation factors. Moreover, a significant fraction of ribosomes bore ribosomal ligands typically associated with translational inactivity: In a subpopulation of 80S ribosomes (13.25%) resolved to 5.64 Å global resolution (Fig. S5A, Fig. S3, Fig. S4C) the decoding center was found to be obstructed with the protein LYAR ^27^, which was consistent with the high abundance of LYAR detected in the 80S fraction by MS (Table S1). Another subpopulation of 80S ribosomes (16.89%) resolved to 4.34 Å global resolution (Fig. S5B, Fig. S3, Fig. S4E) showed P- and E-sites were occupied by the protein IFRD2 ^28^. These combined results demonstrate that ribosomes in the monosome fraction mostly contained idle 80S complexes that were therefore functionally distinct to 80S complexes of the polysome fraction.

**Fig. 2:**
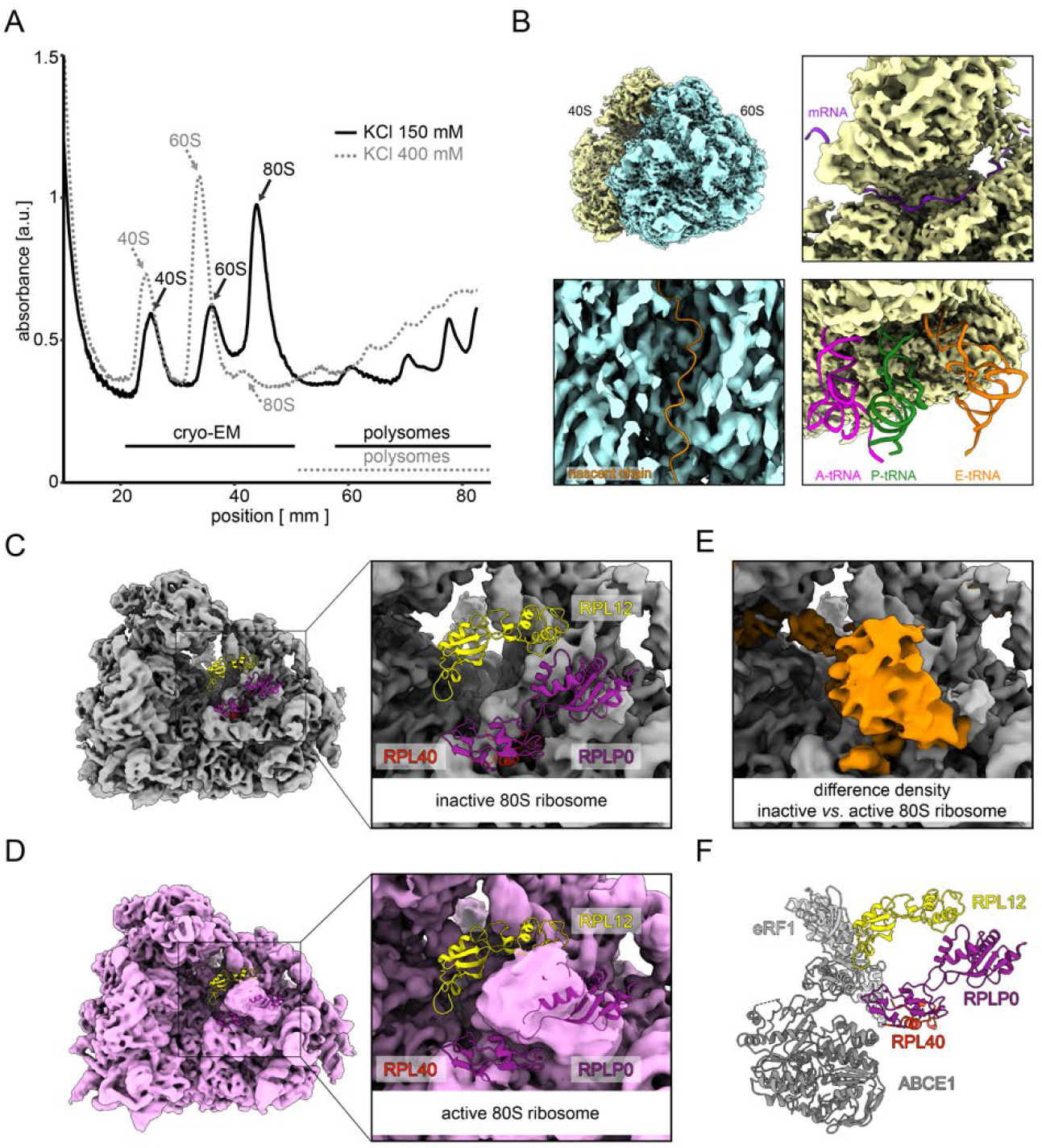
Cryo-EM analysis of non-polysomal ribosomal complexes from MCF7 cells. A) Polysome profiling of MCF7 cells under low and high salt conditions. Ribosomes associated with mRNA remain in their 80S assembly state under high salt, whereas empty 80S ribosomes dissociated into 40S and 60S subunits. Fractions combined and investigated with cryo-electron microscopy are indicated. B) Top left: local-resolution-filtered cryo-EM density of inactive human 80S ribosomes after 3D multi-body refinement. Large (blue; 3.09 Å global resolution) and small (yellow; 3.72 Å global resolution) ribosomal subunits are indicated. 60S, large 60S ribosomal subunit; 40S, small 40S ribosomal subunit. Top right: the mRNA channel is devoid of density for an mRNA molecule (PDB-6YAL)^54^. Bottom left: density of the 60S subunit cut through the polypeptide exit tunnel. The ribosomal exit tunnel site is devoid of density for a nascent polypeptide (PDB-3JAJ)^55^. Bottom right: the intersubunit space is vacant and devoid of density for tRNAs (PDB-5MC6 and PDB-5LZV)^30,56^. C&D) Comparison of the inactive (C) and active human ribosome (D) (EMD-10674)^57^. In both cases, cryo-EM densities are shown at 6-Å resolution for clarity. Atomic models for the components of the P-stalk area have been superposed (PDB-3JAH)^58^. E) Zoomed view of positive difference density (in orange) calculated by the subtraction of the reconstruction shown in C from the reconstruction shown in D. Parts of C colored as previously. F) Positions of eRF1 and ABCE1 relative to RPL40 and the stalk base proteins RPLP0 and RPL12 (PDB-3JAH)^58^.

Interestingly, detailed inspection of the 80S cryo-EM reconstruction indicated increased plasticity of ribosomal proteins for the ribosomal P-stalk area (RPL12, RPL40, RPLP0, RPLP1 and RPLP2, Fig. 2C), as compared to canonical translating ribosomes (Fig. 2D&E)^29^. The ribosomal P-stalk base provides crucial interfaces for binding of translational GTPases and release factors, but also ribosome splitting and recycling factors such as eRF1 and ABCE1 displayed in Fig. 2F ^30^. Higher plasticity of the P-stalk base area may therefore provide a structure-based rationale for the stability of inactive 80S ribosomes by impeding the binding of ribosome splitting factors. Notably, a similar set of ribosomal proteins was previously observed to be specifically ubiquitylated and, thus, destabilized in a similar manner in yeast upon oxidative stress-induced translational halt ^31^, suggesting an evolutionary conserved role for stalk plasticity in the regulation of translational activity. In conclusion, our polysome profiling and cryo-EM analysis showed that the vast majority of ribosomes in the 80S fraction were translationally inactive and did not reside on mRNA, clearly distinguishing them from translationally active, mRNA-bound polysomes.

### Inhibition of Translation Produces Inactive 80S Ribosomes and not Free Subunits

Having observed that ribosomes in the 80S fraction were particularly stable and translationally inactive we aimed to clarify their origin. To investigate if they were derived from previously translating 80S complexes we induced translational arrest with arsenite in MCF7 cells and observed a collapsed polysome fraction and a simultaneous buildup of inactive 80S ribosomes (Fig. 3A). Arsenite is known to inhibit translation initiation via phosphorylation of EIF-2α, thereby preventing loading of the initiator methionine-tRNA complex ^32^. Interestingly, other translation inhibitors with different modes of action, such as harringtonine (stalling translation initiation without interfering with translation elongation or termination ^33–35^) and puromycin (inducing premature termination by peptide chain termination ^36^) also led to a strong accumulation of inactive 80S ribosomes, mirroring earlier, published observations using human or mouse cells ^37,38^. We found that arsenite stress led to a complete collapse of the polysome fraction as soon as ten minutes into the treatment (Fig. S6A), and we used the non-polysomal fractions from this time point for structural characterization by cryo-EM analysis. This confirmed that these 80S ribosomes were translationally inactive and virtually identical to the 80S ribosome without arsenite treatment in terms of overall structure and composition (Fig. S6B, see also Materials and Methods). Consistently, compared to the 80S ribosomes without arsenite treatment, densities for ribosome inactivation factors LYAR and IFRD2 were detected at similar stoichiometries (15.52% and 13.02%, respectively, Fig. S5C&D). The high similarity of idle 80S ribosomes observed under unperturbed conditions and after arsenite treatment indicated that inactive 80S ribosomes in general were derived from previously translating 80S ribosomes.

**Fig. 3:**
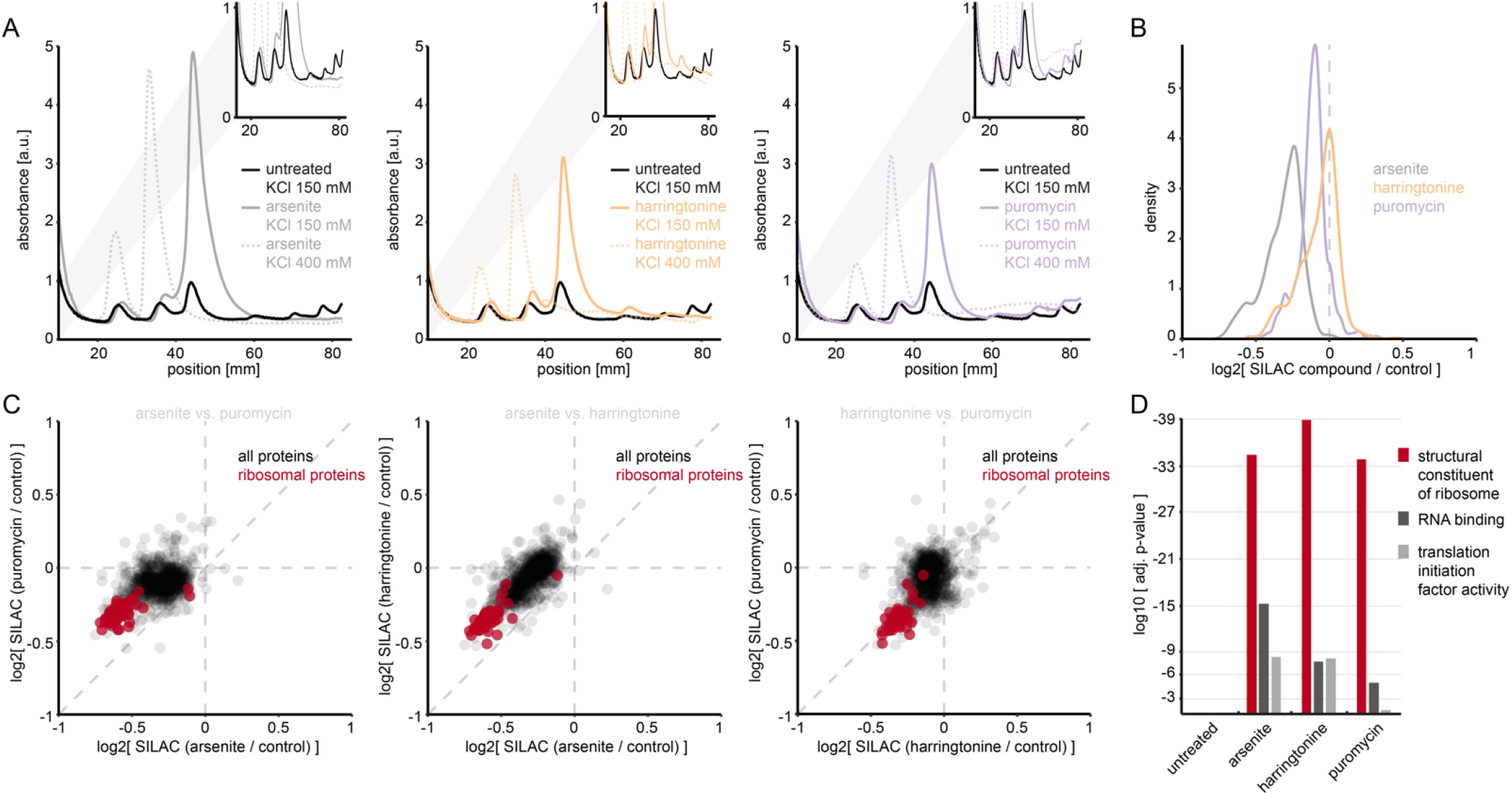
Translation inhibitors expand the pool of inactive 80S ribosomes and induce ribophagy. A) Polysome profiling of MCF7 cells upon translation inhibition. Cells were treated 30 minutes with indicated translation inhibitors and subjected to high and low salt polysome profiling. B) Density plot for expression changes in the total proteome of MCF7 cells upon translation inhibition. Cells of one SILAC label were treated with the indicated translation inhibitor and compared to untreated cells of the complementary SILAC label. Normalization occurred to control samples where both SILAC channels were untreated (for details see Materials and Methods). Shown are means of quadruplicates with label swap filtered for a variance smaller 20 %. D) Scatterplot comparing expression changes in MCF7 cells upon arsenite, harringtonine and puromycin treatment. Same data as in B. Cytosolic ribosomal proteins are highlighted in red. E) Ranked GO enrichment analysis (molecular function) for proteins with decreased expression in MCF7 cells upon translation inhibition. Compared are the top-three enriched terms upon arsenite treatment and their significance levels under different treatment regimens.

### An Increased Pool of Inactive 80S Ribosomes Accelerates the Ribophagic Flux

Recently, we reported that arsenite induces a very fast and drastic form of ribophagy in human cells, eliminating up to 50% of cytosolic ribosomal proteins within 30 minutes ^20^. Since our polysome profiling experiments indicated that translational arrest caused by arsenite, harringtonine and puromycin similarly increased the pool size of idle 80S ribosomes (Fig. 3A), this raised the question if they also triggered the selective elimination of ribosomes. Indeed, SILAC-based quantification of the MCF7 total proteome revealed that all three translation inhibitors induced similar degradation of the same proteins within 30 minutes of treatment (Fig. 3B&C, Table S3). These proteins were highly enriched for gene ontology terms related to the cytosolic ribosome and translation, (p<10E-30 for all three substances, ranked GO enrichment, Fig. 3D), strongly indicative of ribophagy. Thus, our data suggested that translational arrest, irrespective of the agent and its mechanism of action, produced inactive 80S ribosomes, consequently increasing the ribophagic flux. Especially the mechanism of puromycin – leading to unnatural peptide chain termination without natural translation termination ^36^ – prompted us to conclude that the increased availability of inactive 80S complexes and not translation termination triggered increased ribophagy. By this model, the magnitude of the ribophagic flux should be proportional to the strength of translational inhibition. To test this, we challenged MCF7 cells with six increasing concentrations of arsenite and again used SILAC to quantify protein abundances in their total proteome in comparison to untreated cells (Fig. 4A). This revealed a clear dose-response relationship in the ability of arsenite to induce degradation of ribosomal proteins, which was readily reproducible in HeLa cells (Figure S7A). Collectively, we show that the availability of inactive 80S ribosomes and their degradation is a function of translational activity, thereby forming a regulatory connection between protein synthesis and decay.

**Fig. 4:**
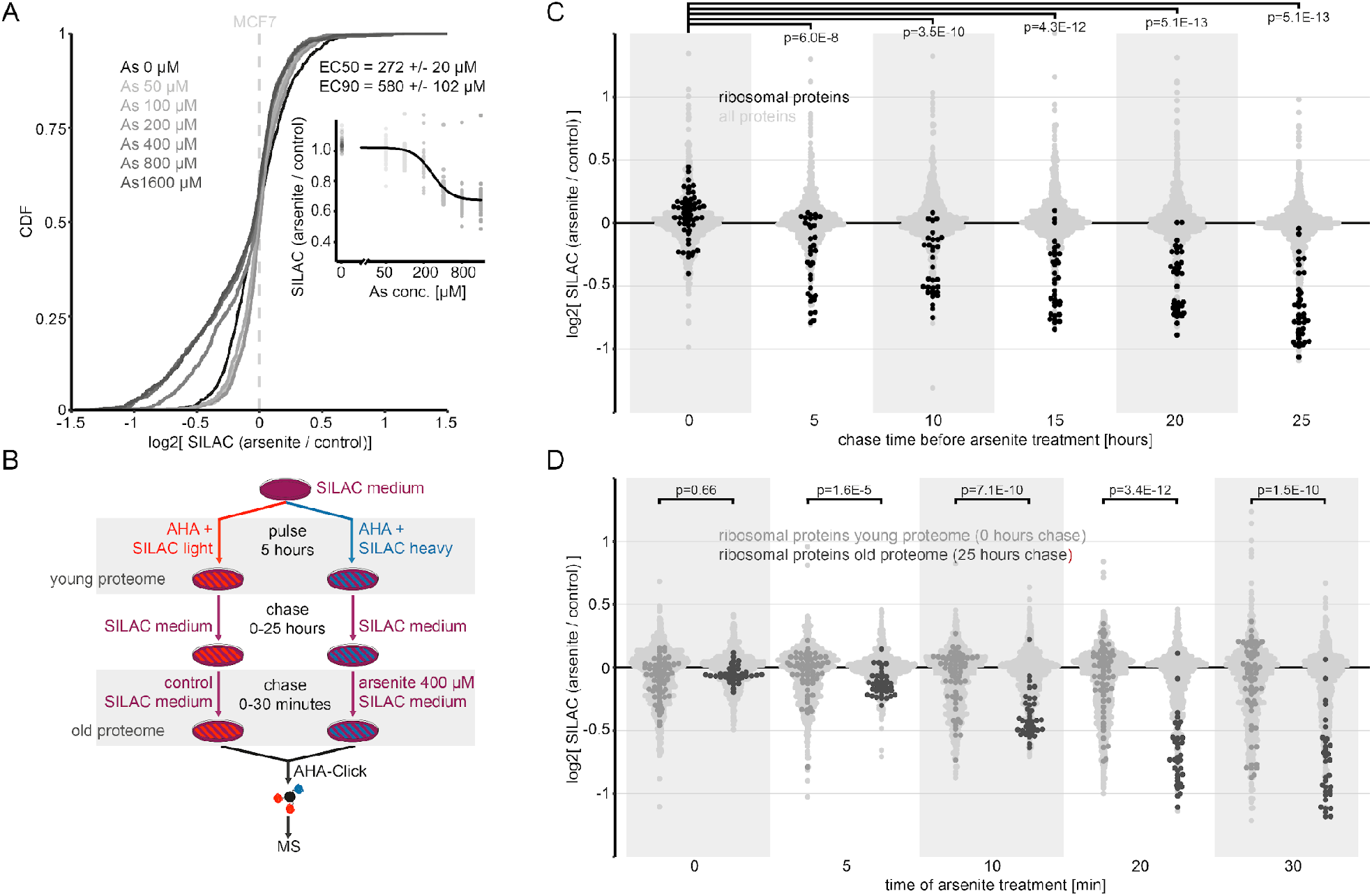
Translational arrest induces degradation of old ribosomes. A) Cumulative distribution of expression changes in the total proteome of MCF7 cells upon increasing doses of arsenite added for 30 minutes of treatment. Insert: Log-logistic model fit against fold-changes of cytosolic ribosomal proteins at different arsenite concentrations. Each dot represents one cytosolic ribosomal protein. B) Proteomic pulse-chase workflow for analysing the effect of protein age on arsenite-induced autophagy. For details see Materials and Methods. C) Dotplot comparing arsenite-induced autophagy within MCF7 proteomes of different ages (5-25 h). Black dots represent cytosolic ribosomal proteins, light grey dots all other identified proteins. Shown are means of duplicate experiments with label swap filtered for a variance smaller 20 %. Pairwise testing occurred for ribosomal proteins with a Wilcox ranksum test between time point 0 hours and each of the other time points as displayed. D) Dotplot comparing arsenite-induced autophagy between the old and new MCF7 proteome directly. Black dots represent cytosolic ribosomal proteins in the old proteome (25 h chase), orange dots cytosolic ribosomal proteins in the new proteome (0 h chase), light grey dots all other identified proteins. Shown are means of duplicate experiments with label swap filtered for a variance smaller 20 %. Pairwise testing occurred with a Wilcox ranksum test between ribosomal proteins of the young and old proteome as displayed.

### Translational Arrest Induces Ribophagic Degradation of Old Ribosomes

Since the ribophagic machinery is assumed to target ribosomes in the 80S assembly state ^18^, and since we had observed that inactive 80S ribosomes were on average much older than other ribosomal assemblies (Fig. 1E), we hypothesized that ribophagy preferentially eliminated old ribosomes. To test this experimentally, we monitored the arsenite-induced degradation of proteins with increasing age. Therefore, we applied azido-homoalanine (AHA) to mark proteomes of different ages (5 hours AHA-labelling pulse followed by 0-25 hours of chase), which were then enriched by click-chemistry for analysis by MS (Fig. 4B). In order to compare protein abundances in the differently aged proteomes of arsenite and untreated cells, this was again combined with SILAC labelling ^39,40^. Indeed, the effect of arsenite on ribosome degradation was strongly dependent on protein age: while young ribosomal proteins (no chase) were nearly untouched by arsenite-induced ribophagy, older ribosomal proteins were progressively degraded with increasing age (Fig. 4C). Direct comparison of a young (no chase) to an old (25 hours chase) proteome over the course of 30 minutes of arsenite treatment emphasized this effect (Fig. 4D). Ribophagy reportedly leads to significant bystander flux, eliminating non-ribosomal, translation-associated proteins, while ingesting ribosomes into autophagic vesicles ^15,20,41^. We also observed bystander flux, however without protein age-dependence (Fig. S7B), indicating that arsenite-induced ribophagy specifically eliminated old ribosomes and not old protein in general. These data demonstrate that arsenite-induced ribophagy accelerates the targeted degradation of 80S ribosomes, thereby eliminating old ribosomal protein, ultimately rejuvenating the ribosomal proteome.

### Constrained Conformational Plasticity of Inactive 80S Ribosomes Upon Translational Halt

In search for a degradation signal on 80S ribosomes we analyzed our differentially treated total proteomes (Fig. 3C) as well as our pulsed-SILAC-AHA timelines (Fig. 4C&D) for arsenite-induced protein modifications using the mass-tolerant search engine MSfragger ^42^. However, we neither found appreciable degrees of known modifications (ubiquitination, phosphorylation, acetylation) nor accumulations of unknown mass adducts (data not shown). In order to understand if there were any other molecular features that cued the degradation of 80S ribosomes upon translational arrest, we revisited our cryo-EM data. Since the overall structures of the inactive 80S ribosomes were very similar in untreated cells and in cells exposed to arsenite for 10 minutes (Fig. 2, Fig. S6) we explored whether their structural plasticity differed, especially with respect to the large-scale, inter-subunit rearrangements that ribosomes undergo during each cycle of translation elongation ^29^. By sorting the 80S complexes according to their 40S-60S rotation we observed that untreated 80S ribosomes exhibited heterogeneous 80S conformational states, with the post-translocation (POST)-like state constituting the most abundant class occurring in 38.16 % of cases (Fig. S7C&D). In contrast, after arsenite exposure inactive ribosomes were conformationally highly homogenous, adopting the POST-like conformational state in 90.55 % of cases (Fig. S7E&F). This indicated that a thus far unknown mechanism limited the conformational plasticity of 80S ribosomes upon arsenite-induced translational arrest where ribosomes mostly maintained the energetically stable POST-like configuration ^43^.

Overall, our structural analysis suggests that the ribophagic flux might not only be controlled by the concentration of inactive 80S ribosomes but also by their conformational state.

## Discussion

### Per-Peptide Normalization Allows for Protein Turnover Studies in Purified Protein Complexes

Determination of protein half-lives in cells that grow and divide has been a long-standing problem, revolving around the conundrum that data generated from systems in disequilibrium do not formally qualify to be fit to a model that requires a steady state, as it is the case for the first-order decay model commonly used to describe protein decay ^23,24,44,45^. In order to still approximate protein turnover, previous studies fitted data from dividing cells to the model, frequently applying a *post hoc* correction factor for cell division ^23,24,44^. Here we introduced PSN to take an alternative approach by first applying a per-peptide normalization, strongly enhancing the fitting process by improving continuity of the data and transforming it into the equilibrium state required by the model. This is conceptually different to previous studies that neglected the equilibrium requirement before fitting. Consequently, PSN allowed us to derive turnover kinetics for ribosomal proteins in four states (free protein, part of free 40S/60S subunit, idle 80S ribosome, or actively translating 80S ribosome), demonstrating that the half-life of the same protein can differ depending on the complex it participates in. Our findings confirm previous work that found many half-lives in the total proteome are better explained by a two-state model for protein decay ^45^, which in the case of ribosomal proteins we can in fact extend to a superposition of at least four distinct decay processes. We note here that data normalized with PSN can of course be fitted to higher-order models, which will undoubtedly benefit from the improved data continuity just as much. PSN rationalizes protein half-life measurements in dividing cells accepting that absolute half-lives will be slightly skewed to the benefit of much improved fitting, which we argue is better than skewing half-lives by fitting non-steady-state protein decay curves to a steady-state model. Thereby, PSN yields the distinctive power to compare protein half-lives between conditions or within different cellular preparations, as evidenced by our analysis of half-lives within polysome profiling fractions. We envision future applications of PSN in studying the proteome dynamics within other complexes (e.g. using immunoprecipitation, size exclusion chromatography etc.), in organelles, or in other non-steady-state scenarios (e.g. drug treatments, development, infection etc.).

### Distinct Pools of Ribosomal Subunits Organize the Ribosome Life cycle

In the canonical view on translation in eukaryotes, 80S splitting is seen as the last step in the termination process, releasing free subunits from the mRNA into the cytosol ^9^. Several of our observations challenged this model, instead indicating that 80S complexes reunite into stable, inactive 80S complexes after translation termination (Fig. 5): First, inhibition of translation with various drugs consistently led to the accumulation of idle 80S ribosomes (Fig. 3A). Second, these inactive 80S ribosomes were structurally identical to inactive 80S ribosomes from untreated cells (Fig. 2B & Fig. S6B). Third, protein half-lives showed that ribosomal proteins were strongly stabilized in the 80S or polysome fraction, while this was not the case in the 40S or 60S fraction (Fig. 1E), suggesting the existence of distinct subunit pools that did not intermix. Importantly, all three observations are in direct agreement, where the formation of inactive 80S complexes after translation provides a mechanism to avoid mixing with nascent, free 40S and 60S subunits. Based on these observations we propose that inactive 80S complexes are a normal transient state arising from translating ribosomes that have finished one round of translation, potentially awaiting to engage in a new round (Fig. 5). Indeed, it has been shown that recycling of 80S complexes can be linked to translation initiation via ABCE1 – a central component of the ribosome splitting apparatus.^46^ Moreover, a mechanistic study on translation initiation from idle 80S ribosomes found in an *in vitro* translation system with human components that formation of the 48S initiation complex occurred equally well from inactive 80S assemblies or free 40S subunits in the presence of the human splitting apparatus (PELO-HBS1L-ABCE1)^47^. Additionally, in yeast it was reported that upon nutrient withdrawal inactive 80S ribosomes accumulate and require the yeast ribosome splitting apparatus (Dom34-Hbs1) to resume translation when nutrients are replenished ^48^. This indicates that inactive 80S ribosomes are stable yet biologically accessible entities. Our cryo-EM analysis suggested that inactive ribosomes within the 80S fraction might be splitting-incompetent because of the elevated plasticity in their binding sites for ribosome splitting and recycling factors (Fig. 2C-F). In order to regain the ability to split and engage in translation initiation (Fig. 5) idle 80S complexes would therefore require reconstitution of the p-stalk, offering a regulatory mechanism by which inactive 80S ribosomes might become reactivated towards the pool of actively translating ribosomes. Indeed, in yeast it has been shown that the flexible ribosomal stalk plays an important role in translation regulation ^49^ and that after translational arrest inactive 80S ribosomes return to normal function ^48^.

**Fig. 5:**
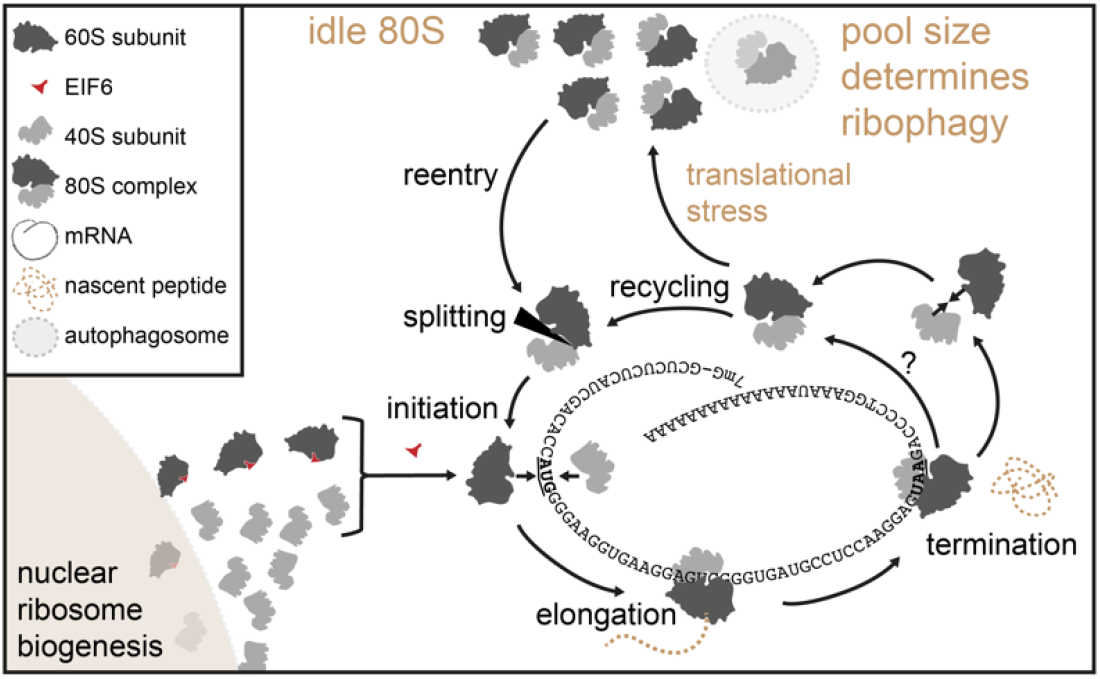
Schematic representation of the ribosome lifecycle in human cells. Translation-competent ribosomal subunits are stored in inactive 80S ribosomes after translation to keep them separate from nascent subunits carrying anti-association factors.

An open question is how subunits reunite after being released from the mRNA. This could occur spontaneously because the 40S and 60S subunits have high affinity towards each other and are in close proximity after splitting with very limited diffusion rates due to the high degree of molecular crowding in the cytoplasm ^50^. In fact, it is known that eIF6 binds to nascent 60S subunits to prevent their spontaneous joining with 40S subunits ^51^. In accordance with this, we found eIF6 at a 1:1 ratio relative to large subunit proteins in the 60S fraction (Table S1) and our cryo-EM analysis similarly indicated that eIF6 was present on 30 % of the identified 60S subunits (Fig. S8, Fig. S3, Fig. S4K). In contrast, all other fractions contained eIF6 only at a ratio of around 1:100 (Table S1). This implies that the free 60S complexes bear eIF6 and are therefore nascent ^51^, whereas mature 60S subunits in idle or translating 80S complexes do not bear EIF6 and therefore spontaneously rejoin into 80S complexes after splitting. Notably, this is in line with our half-life measurements where we observed free 60S subunits to be very young (Fig. 1E). Considering that translating 80S complexes are often in the proximity of other 80S complexes on the same mRNA or on the surface of the endoplasmic reticulum, re-joining could also occur between distinct 80S complexes thereby exchanging subunit partners. Another, yet, less-documented possibility is that inactive 80S ribosomes arise by a drop-off mechanism, releasing intact 80S complexes from mRNA during premature termination.^52^

Conceptually, a pool of 80S ribosomes distinct from free subunits comes with a number of advantages. First, functional 80S ribosomes are kept apart from potentially immature or dysfunctional, nascent subunits. Second, if nascent subunits were always mixed with old subunits, all subunits would have the same likelihood for being degraded, and destroying only just generated subunits by chance cannot be energetically favorable. Third, in case the protein synthesis apparatus is failing and translation halts, affected 80S ribosomes can be removed in a targeted and potentially conformation-selective fashion through ribophagy, allowing free nascent subunits to replace them. Collectively, a pool of 80S ribosomes distinct from free subunits should ensure a functionally tested and quantitatively attuned translational apparatus at all times.

### Translational Activity Regulates Ribophagic Flux

Currently ribophagy is seen as a mechanism that is initiated by nutrient deprivation or mTOR inhibition to liberate the significant fraction of amino acids and nucleotides stored in ribosomes ^14,15,17,18,48^. Indeed, starvation experiments in human cells saw clear enrichment for ribosomal and translation-associated proteins eliminated via autophagy over the course of 24 hours ^15^. Both starvation and mTOR inhibition are unspecific ways for the induction of autophagy, raising the question in how far ribophagy is any different to general autophagy. Our data shows that ribophagy can be a distinct process acting on the time scale of minutes, when translation is directly inhibited. This rapid elimination of ribosomes is markedly different to the long term autophagic degradation of ribosomes upon mTOR inhibition or starvation, which occur on the time scale of several hours to days ^15,17^. We propose that ribophagy is a substrate-driven process, where an increased 80S pool offers more chances to the autophagic machinery to degrade mature ribosomes. This means that ribophagy can be defined as a distinct autophagic process that reacts to the accumulation idle 80S ribosomes. Under this definition, the actual ribophagic flux is determined by the availability of inactive 80S ribosomes and not by the activation status of the autophagy machinery itself. Interestingly, classic stimulation of autophagy via nutrient withdrawal or mTOR inhibition also lead to translation inhibition and a dramatic increase in inactive 80S ribosomes ^53^, clarifying why ribophagy is seen as a subprocess of general autophagy. Our structural data indicates that an accumulation of inactive ribosomes in the POST-like state may flag 80S ribosomes for ribophagy, but whether and how their reduced conformational plasticity could serve as a degradation signal remains to be investigated. In summary, we propose that ribophagy is a normal subprocess of general autophagy during starvation, however, it is substrate-driven and can occur independently when translation halts and idle 80S ribosomes accumulate.

## Acknowledgments

We thank Jana Zecha (TU Munich, Germany) and Bernhard Küster (TU Munich, Germany) for help with implementing TMT-SILAC in MaxQuant. We thank Gregor Mönke (EMBL Heidelberg, Germany) for discussion and help on benchmarking PSN. We acknowledge the data storage service SDS@hd and bwHPC supported by the Ministry of Science, Research and the Arts Baden-Württemberg, as well as the German Research Foundation (INST 35/1314-1 FUGG and INST 35/1134-1 FUGG). We also acknowledge access to the infrastructure of the Cryo-EM Network at the Heidelberg University (HDcryoNET). This work was supported in part by the Excellence Cluster CellNetworks (to JK) and by an ERC Advanced grant ‘Transfold’ (to BB).

## Author contributions

Conceptualization: JT, JK; Methodology: JT, MA, MB, MJ; Investigation: JT, MA, MB, MJ; Visualization: JT, MA, MJ; Funding acquisition: SP, BB, JK; Project administration: JT, JK; Supervision: GK, SP, BB, JK; Writing – original draft: JT, MA, SP, JK; Writing – review & editing: JT, MA, MB, MJ, GK, SP, BB, JK

All authors declare that they have no competing interests.

## Data availability

Data is currently being uploaded to ProteomeXchange and EMDB.

## Supplementary Materials

Figs. S1: Proteomic quantification of MCF7 polysome profiling fraction and comparison of normalization procedures for TMT-SILAC data.

Figs. S2: Benchmarking peptide sum normalization.

Figs. S3: Cryo-EM processing workflow for ribosomal complexes with and without arsenite treatment.

Figs. S4: Local resolution estimation and Fourier shell correlation curves.

Figs. S5: Identification of LYAR and IFRD2 bound to translationally inactive 80S ribosomes without and with arsenite treatment.

Figs. S6: Cryo-EM structure of the inactive human 80S ribosome after arsenite treatment.

Figs. S7: Arsenite treatment constrains conformational plasticity of the inactive 80S ribosome.

Figs. S8: Identification of eIF6 bound to 60S ribosomal subunits without arsenite treatment.

Table S1: Label-free quantification of proteins in MCF7 polysome profiling fractions.

Table S2: Protein half-lives from the MCF7 total proteome and MCF7 polysome profiling fractions.

Table S3: Changes in the MCF7 total proteome upon translation inhibition.

## Materials and Methods

### Cell Culture and SILAC

The cell lines MCF7 (ATCC, RRID:CVCL_0031) and HeLa (ATCC, RRID:CVCL_0030) were maintained in Dulbecco’s Modified Eagle’s Medium (DMEM) for SILAC supplemented with 10% dialysed FBS (Gibco 26400-044) and Pen-Strep (100 U / ml penicillin, 100 mg / ml streptomycin, Gibco 15140-122) at 37 °C, 5 % CO_2_. DMEM for SILAC (Silantes 280001300) was supplemented with 1 mM L-lysine and 0.5 mM L-arginine of the individual SILAC labels (Silantes 211104113, 201204102, 211604102, 201604102) as well as 1.7 mM light L-proline and 1 x GlutaMAX (Gibco 35050061). For full labelling the intermediate or heavy SILAC label was introduced during six passages in intermediate or heavy DMEM for SILAC, respectively. All experiments were performed on MCF7 cells, except when confirming the dose-response of arsenite treatment on HeLa cells (Fig. S5B). Cells were seeded at a density so that they reached 70 % confluence after three days of expansion in culture (1 million MCF7 cells for a 15 cm culture dish). Except for the protein half-life experiments, all experiments were performed three days after seeding. The cell culture timing for the protein half-life experiments is detailed below.

### Purification of Total Proteomes for Proteomics Using Single-Pot-Solid-Phase-Enhanced-Sample-Preparation (SP3)

For proteomic sample preparation of total proteomes a modified version of the SP3 protocol was used ^59^. Cell pellets or lysates were topped off to a total volume of 900 µl with SP3 lysis buffer (tris-Cl 50 mM, SDS 0.05 %, DTT 10 mM). Lysis was facilitated by pipetting and vigorous vortexing. Samples were reduced at 90 °C, 700 rpm shaking, for 30 minutes and then cooled on ice. Twenty µl CAA 1 M was added along with 1 µl benzonase (Novagen 70664) and samples incubated 2 hours at 37 °C, 700 rpm shaking. One hundred µl denaturation solution (EDTA 200 mM, SDS 10 %) was added and samples vigorously vortexed. Four hundred µl SP3 (GE 44152105050250) beads were preconditioned by washing with 1 ml MilliQ water 3 times, before reconstitution in 1 ml MilliQ water. Twenty µl preconditioned SP3 beads were mixed to the samples by vortexing. One ml acetonitrile was added and binding of proteins occurred for 15 minutes at room temperature. Beads were captured on a magnetic rack, supernatants discarded and beads washed three times with 2 ml EtOH 70%, while attached to the magnet. Tubes were spun down and put back on the magnet in order to remove all residual EtOH. Beads were taken up in 100 µl TEAB 20 mM (Sigma T7408) containing 500 ng trypsin/LysC (Promega V5073) and digested for 16 hours at 37 °C, 700 rpm. Subsequently, peptides were cleaned up using an Oasis PRiME HKB µElution Plate. Peptides were taken up in 15 µl formic acid 1 % before analysis by HPLC-MS with the method detailed in each specific experimental section or 80 µl high pH buffer (NH_4_COOH 20 mM) for fractionation at high pH, or 20 µl TEAB 20 mM for TMT-labelling, as outlined below.

### Purification of Ribosomal Complexes by Polysome Profiling and an SP3 Modification for the Proteomic Sample Preparation from Sucrose Gradient Fractions

For one polysome profiling experiment approx. 10 million MCF7 cells grown on a 15 cm dish were used. The media was discarded and residual media removed by tapping the culture dish onto a paper towel. Cells were rinsed with 10 ml ice-cold harvest buffer (MgCl_2_ 10 mM, cycloheximide (Santa Cruz sc-3508A) 100 µg/ml in PBS), which was subsequently discarded and again residues removed by tapping the culture dish onto a paper towel. The dish was transferred onto ice and cells scraped into 100 µl ice-cold lysis buffer 5 x (NP40 5 %, tris-Cl 250 mM, MgCl_2_ 50 mM, KCl 700 mM, cycloheximide 500 µg/ml, DTT 5 mM, 1 x EDTA-free protease inhibitor (Roche 11873580001), to 100 µl add 15 µl DNase I and 1.5 µl RNASin Plus (Promega) before use), which becomes diluted to 1x with residual PBS on the dish surface.^60^ Lysates were transferred to a fresh tube, vortexed and incubated on ice for 10 minutes before passing through a 26-gauge needle 10 times in order to shear genomic DNA. Consequently, samples were cleared by centrifugation with 20000 g at 4 °C for 5 minutes. Sucrose gradients of 5-45 % were prepared in sucrose buffer (tris-Cl 50 mM, MgCl_2_ 10 mM, KCl 140 mM or 400 mM, 1 x EDTA-free protease inhibitor, cycloheximide 100 µg/ml) on a BIOCOMP153 gradient station (BioComp Instruments). Supernatants were transferred onto the sucrose gradients and subjected to 3.5 hours of ultracentrifugation with 35000 rpm in a Sorvall WX90 ultracentrifuge (Beckman) and the rotor SW40Ti. Gradients were then collected into 60 fractions using a BIOCOMP153 gradient station. Subfractions representing the 40S, 60S, 80S and polysome fractions were combined according to the UV trace. For one experiment series identical subfractions were combined, however, the exact fraction numbers varied slightly between experiment series and were adapted accordingly.

As the sucrose gradient in the polysome profiling contained KCl, which precipitates SDS, we used a modification of the SP3 protocol presented above. The total volume of the combined fractions amounted to approximately 1 ml for the 40S, 60S and 80S fractions, of which 900 µl were transferred to a 2 ml tube. Ten µl DTT 1 M was added and samples were reduced at 90 °C, 700 rpm shaking, for 30 minutes and then cooled on ice. Twenty µl CAA 1 M was added along with 1 µl benzonase and samples incubated 2 hours at 37 °C, 700 rpm shaking. One hundred µl denaturation solution (EDTA 200 mM, 20 M guanidinium chloride) was added and samples vigorously vortexed. Four hundred µl SP3 beads were preconditioned by washing with MilliQ water 3 times, before reconstitution in 1 ml MilliQ water. Twenty µl preconditioned SP3 beads were mixed to the samples by vortexing, before addition of 1 ml acetonitrile. Binding of proteins occurred for 15 minutes at room temperature. Beads were captured on a magnetic rack, supernatants discarded and beads washed three times with 2 ml EtOH 70%, while attached to the magnet. An additional round of washing was performed were beads were taken off the magnet and disintegrated into 1 ml EtOH 70 %. Samples were put back onto the magnet in order to remove all EtOH. Beads were taken up in 100 µl TEAB 20 mM containing 500 ng trypsin/LysC and digested for 16 hours at 37 °C, 700 rpm and peptides cleaned up using a Oasis PRiME HKB µElution Plate. Peptides were taken up in 15 µl formic acid 1 % before analysis by HPLC-MS with the method detailed in each specific experimental section or 20 µl TEAB 20 mM for TMT-labelling. An exception was the polysome fraction, where the combined volume of the subfractions was approximately 4 ml. These 4 ml were collected in a 15 ml falcon tube and treated analogously to the other fractions. Volumes were adjusted except for benzonase and SP3 beads, for which volumes were kept the same. After collecting the SP3 beads on a 15 ml magnetic rack and discarding the supernatant, beads were taken up in two times 1 ml EtOH 70 % in order to be transferred to a fresh 2 ml tube. From that point on samples were treated identically to the other fractions.

### Determination of Protein Half-Lives by TMT-SILAC

For all protein half-life measurements two replicates were generated from MCF7 cells that included a SILAC label swap. Explicitly, one replicate was produced from SILAC heavy cells switched to light SILAC media and another replicate from SILAC light cells switched to heavy SILAC media. For determining half-lives of the total proteome seven timepoints were generated. In the case of the total proteome 1 million MCF7 cells of one SILAC label were seeded on 15 cm dishes and expanded for 3 days. Subsequently, the media was discarded, cells were washed twice with PBS to remove residual media before addition of new media with the complementary SILAC label. Cells were harvested after additional 0, 1, 2, 4, 8, 16 and 32 hours of culture after switching the SILAC media. For the harvest, media was discarded, cells put on ice and scraped into 10 ml ice-cold PBS and transferred to a falcon tube. Residual cells were scraped into another 10 ml ice-cold PBS, combined with the rest and spun down for 5 minutes with 1000 g at 4 °C. Cells were lysed in 1 ml SP3 lysis buffer (tris-Cl 50 mM, SDS 0.05 %, DTT 10 mM) by vigorous pipetting and vortexing before 200 µl of the lysate was used for further purification by conventional SP3 as outlined above.

For determining half-lives of proteins in polysome fractions again two replicates with label-swap were generated. However, in order to increase the resolution of our analysis nine timepoints were taken. As the quality of polysome profiling suffers if samples are frozen, all cells were harvested at the same time. This required that the SILAC media was changed at appropriate time distances towards one common harvest point. Therefore, 1.3 million MCF7 cells of one SILAC label were seeded onto 15 cm dishes and harvested after a total of 4 days in culture. Cells were washed twice with PBS and switched to the complementary SILAC label 0, 2, 4, 8, 12, 16, 20, 24 and 32 hours before those 4 days had passed. Cell harvest, polysome profiling and protein clean-up occurred as described above. In the case of the total proteome the first 2-7 TMT channels were occupied by the time points 1-32 hours, the TMT channel 8 and 10 by replicates of the infinity time point (for the replicate starting from SILAC light cells the infinity timepoint were SILAC heavy cells and *vice versa*) and the channels 1 and 9 by replicates of the time point 0. Replicate channels were produced from the same peptides in order to assess the reproducibility within one experiment. The reproducibility was found very good for all replicates so that for the polysome profiling experiments all channels were used for time points, i.e. for polysome profiling fractions the time points 0-32 hours were allocated to the TMT channels 1-9 and the infinity time point to TMT channel 10.

In order to label identical amounts of peptides for each TMT channel, peptide concentrations were assessed using a Quantitative Colorimetric Peptide Assay (Pierce 23275) on 10 % of each sample. The amount of peptides that were labelled was oriented on the sample with the lowest peptide concentration in one particular experiment series. In the case of the total proteome 10 µg peptides per time point were used, in the case of the polysome fractions 5 µg of peptides were used. Identical amounts of peptides for each timepoint were adjusted to a total volume of 20 µl in TEAB 50 mM. Consequently, 5 µl of TMT labelling reagent in acetonitrile (Thermo 90113, see manufacturer manual for dilution) was added and labelling allowed to occur for 1 hour at room temperature. Labelling was quenched by addition of 1.5 µl hydroxylamine 5 % and another 15 minutes of incubation. Samples for all 10 TMT channels of one experiment were combined, completely dried by SpeedVac and taken up in 80 µl high-pH buffer (NH_4_COOH 20 mM) for fractionation at high pH as described below. In the case of the total proteome replicates were analysed in 16 fractions. Therefore, the first 8 of the 40 collected fractions were discarded and the following combined to 16 fractions using the scheme 1+17/…/16+32. In the case of the polysome profiling experiments, samples were analyzed in 8 fractions. Therefore, the first 8 of the 40 collected fractions were discarded and the following combined to 8 fractions using the scheme 1+9+17+25/…/8+16+24+32.

Fractions were again dried down completely and taken up in 15 µl formic acid 1 % before analysis by MS. HPLC-MS occurred on a Fusion MS using the 2 hour gradient described below and the standard SPS-MS3 method for TMT recommended by the manufacturer.

### Azidohomoalanine (AHA) Labelling for the Extraction of Proteomes of Different Ages

Briefly, in order to monitor the effect of arsenite on MCF7 sub-proteomes that were aged 0-25 hours, intermediate SILAC-labelled MCF7 cells were pulsed for five hours with either SILAC heavy or SILAC light labels, respectively, in the presence of AHA (Jena Bioscience CLK-AA005). Subsequently, this ‘young’ proteome was allowed to age by chasing cells with intermediate SILAC media without AHA for 0-25 hours at five-hour intervals. Heavy-labelled cells from each age group were then treated with arsenite (Santa Cruz sc-301816) for 30 minutes to induce autophagy, which were then combined with light-labelled untreated cells, and subjected to click-chemistry to selectively isolate the aged AHA-labelled proteomes. Finally, upon MS the heavy-over-light SILAC ratio was used to determine arsenite-induced effects on the proteome for each age group, whereas background signal was relegated to the intermediate SILAC channel.

Explicitly, cells were first subjected to AHA-labelling. Therefore, 1.4 million SILAC intermediate MCF7 cells were expanded on 15 cm dishes for 3 days in standard SILAC intermediate DMEM (see Mammalian Cell Culture and Stable Cell Lines). The media was discarded, cells washed twice with PBS and depletion media (reduced component DMEM (AthenaES 0430), sodium bicarbonate 3.7 g/l, Sodium Pyruvate 1 mM, HEPES 10 mM, GlutaMax 1 x, L-proline 300 mg/l, L-cystine 63 mg/L, L-leucine 105 mg/l, dialyzed FBS 10 %) added for 1 hour of incubation at normal culture conditions. The depletion media was discarded and replaced by labelling media (depletion media supplemented with L-lysine 146 mg/l, L-arginine 84 mg/l, L-azidohomoalanine 18.1 mg/l) for 5 hours of incubation at normal culture conditions. Note that the SILAC heavy and light labels were introduced via the labelling media and during the 5 hours of AHA-labelling. Subsequently, cells were washed twice with PBS and incubated in standard SILAC intermediate DMEM for 0-25 hours of label chase. Following the chase period AHA-SILAC-heavy cells were treated with 400 µM arsenite for 0-30 minutes, while AHA-SILAC-light cells were left untreated. Simultaneously a second replicate was produced with SILAC label swap. The media was discarded and cells were transferred onto ice. Cells were harvested into 10 ml ice-cold PBS by scraping and transferred to a falcon tube. Residual cells were scraped into another 10 ml, which were combined with the first, and spun down with 1000 g for 5 minutes at 4 °C. Supernatants were discarded and AHA-labelled proteins enriched using the Click-it Protein Enrichment kit (Invitrogen C10416) according to the instructions of the manufacturer. Proteins were eluted from the beads into 200 µl digestion buffer (tris-Cl 100 mM, acetonitrile 5 %, CaCl_2_ 2 mM) containing 500 ng trypsin/LysC during 16 hours incubation at 37 °C, 1000 rpm shaking in 2 ml tubes. Peptides were cleaned up using an Oasis PRiME HKB µElution Plate and taken up in 15 µl formic acid 1 % before analysis by HPLC-MS on a QExactive HF with the standard method outlined below.

### Quantification of Ribophagy Upon Translation Inhibition

During conventional SILAC experiments for the effect of arsenite-treated against untreated cells we observed that unnormalized ratios were always more extreme than the ratios presented by MaxQuant’s standard SILAC normalization, independent of the SILAC channels used for the comparison. This indicated that most of the cytosolic proteome was affected by bystander autophagy, so that normalization to the median ratio between SILAC channels underestimated the arsenite-induced degradation effect. To capture this effect and still be able to correct for small differences in the number of cells used for each SILAC channel we took two measures: First, we started out from a single population of SILAC intermediate-labelled MCF7 cells (Lys4, Arg6), of which we seeded 0.2 million cells onto 3.5 cm dishes and expanded them in either SILAC heavy (Lys8, Arg10) or SILAC light (Lys0, Arg0) media for 3 days. Second, we combined quadruplicates of heavy and light labelled cells, which were both left untreated in order to define the exact difference between the two SILAC channels. The mean ratio for each of the proteins detected in these samples was used as a correction factor to normalize for mixing errors in the treated samples. Treatment occurred for 30 minutes with 400 µM arsenite, 200 µM puromycin (Gibco A1113803) or 20 µM harringtonine (Sigma SML1091), respectively. Treatments occurred on quadruplicates with label-swap, i.e. two dishes of light cells were treated and compared to two dishes of untreated heavy cells, and *vice versa* for the label swap. Total proteomes were extracted by SP3 as described above and samples analyzed on 2-hour gradients by HPLC-MS on a QExactive HF with the standard method outlined below.

### High pH Reversed-Phase Fractionation of Proteomic Samples

Fractionation at high pH occurred on an Agilent Infinity 1260 LC system (Agilent) using a Phenomenex Gemini 3 µM C18, 100 × 1 mm column (Phenomonex). Buffer A was NH4COOH 20 mM, buffer B was 100 % acetonitrile. The following gradient was used for all applications described in this manuscript: 0-2 minutes 0 % B, 2-60 minutes linear gradient to 65 % B, 61-62 minutes linear gradient to 85 % B, 62-67 minutes 85 % B, 67-85 minutes 0 % B. Eluates were collected in 40 fractions and combined as described in the individual paragraphs. The initial 8 fractions up to approximately 18 % B contained in samples extracted polysome fractionation RNA contaminations and were therefore discarded. For consistency this occurred for all other samples as well.

### HPLC-MS for the Detection and Quantification of Proteomic Samples

Separation by HPLC prior to MS occurred on an Easy-nLC1200 system (Thermo Scientific) using an Acclaim PepMap RSCL 2 µM C18, 75 µm x 50 cm column (Thermo Scientific) heated to 45 °C with a MonoSLEEVE column oven (Analytical Sales and Services). Buffer A was 0.1 % formic acid, buffer B was 0.1 % formic acid in 80 % acetonitrile. The following gradient was used for all applications described in this manuscript: 0 minutes 3% B, 0-4 minutes linear gradient to 8 % B, 4-6 minutes linear gradient to 10 % B, 6-74 minutes linear gradient to 32 % B, 74-86 minutes linear gradient to 50 % B, 86-87 minutes linear gradient to 100 % B, 87-94 minutes 100 % B, 94-95 linear gradient to 3 % B, 95-105 minutes 3 % B.

MS detection occurred on either a QExactive HF or Fusion mass spectrometer (Thermo Scientific) with the method specified in the experimental outlines above. Standard method for the QExactive HF was MS1 detection at 120000 resolution, AGC target 3E6, maximal injection time 32 ms and a scan range of 350-1500 DA. MS2 occurred with stepped NCE 26 and detection in top20 mode with an isolation window of 2 Da, AGC target 1E5 and maximal injection time of 50 ms. Standard method for the Fusion was MS1 detection in orbitrap mode at 60000 resolution, AGC target 1E6, maximal injection time 50 ms and a scan range of 375-1500 DA. MS2 detection occurred with an HCD collision energy of 33 in ion trap top20 mode with an isolation window of 1.6 Da, AGC target 1E4 and maximal injection time of 50 ms.

### MS Database Search

All MS raw files were searched using MaxQuant (1.6.0.16) ^61^. The database searched was the reviewed UniProt human proteome (search term: ‘reviewed:yes AND proteome:up000005640’, 20216 entries, retrieved 11 September 2017) and the default Andromeda list of contaminants. All settings were used at their default value, except for specifying SILAC configurations and indicating the appropriate number of fractions per sample. For the label-free quantification of ribosomal proteins in polysome profiling fractions, for the quantification of new and old proteins in MCF7 sub-proteomes purified with AHA-labelling, and for the quantification of ribophagy upon translation inhibition the ‘match-between-runs’ option was activated. For the quantification of new and old proteins in MCF7 sub-proteomes purified with AHA-labelling, additionally, the ‘requantify’ option was activated. For the label-free quantification of ribosomal proteins in polysome profiling fractions iBAQ quantification was activated. TMT-SILAC data was searched with the parameters adapted from Zecha et al. ^24^. TMT isotope impurities were specified in Andromeda according to the information provided by the manufacturer. In the type section reporter ion MS3 and TMT 10plex was selected and reporter mass tolerance set to 0.01 Da. Lysine 8 and Arginine 10 were defined in Andromeda as variable modifications and subsequently selected as such in the MaxQuant modifications section next to Oxidation and N-terminal acetylation. Peptide mass tolerance in the instrument section was set to 5 ppm. In the identification section the minimum score for modified peptides and the minimum delta score for modified peptides were set to 0. In the MS/MS – ITMS section the MS/MS – ITMS match tolerance was set to 0.4 Da and water loss was deselected. All other settings were used at their default value.

### Processing and Analysis of TMT-SILAC Data

Data was processed according to Zecha et al. with additional peptide sum normalization. Therefore, data was retrieved from the ‘evidence’ table provided by MaxQuant. PSMs in the evidence table were filtered to remove ‘Potential contaminants’ and ‘Reverse’ matches to the decoy database. Because of their uncertain TMT labelling status PSMs of acetylated N-terminal peptides (‘Acetyl’ in the ‘Modification’ column) were removed as well. For all following steps corrected reporter ion intensities were used (‘Reporter intensity corrected’).

Any PSM in the cleaned-up evidence table containing a lysine 8 or arginine 10 modification was assigned to the SILAC heavy dataset, whereas any PSM that did not carry any such modification was assigned to the SILAC light dataset. PSMs were collapsed into peptides by summing up the reporter ion intensities for each TMT channel from all PSMs of a peptide. The two resulting tables contained for each peptide ten reporter ion intensities, which quantified synthesis or degradation (depending on SILAC heavy or light). Additionally, each peptide was annotated with the uniprot identifier (‘Leading razor protein’) so that peptides from the same host protein could be merged later on. Peptide sum normalization (PSN) can only be applied when TMT peptide intensities for both SILAC channels are available. The two distinct tables of SILAC light and heavy TMT peptide intensities were therefore merged by peptides, discarding peptides only quantified in one SILAC channel. Absolute MS3 reporter intensities depend on the MS1 signal, i.e. the combined SILAC intensity of all time points, which can differ greatly between SILAC channels. Because they are derived from cells carrying only one SILAC label TMT peptide intensities should be identical between time point 0 of the degradation SILAC channel, and time point infinity of the synthesis SILAC channel. This can be used to adjust TMT peptide intensities from either channel to the same baseline intensity. For each peptide in each SILAC channel all TMT reporter intensities were multiplied by a common factor to align time points 0 and infinity:

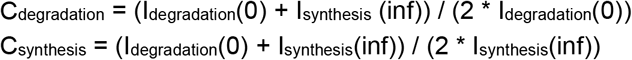

Where C_degradation_ is the correction factor for one peptide in the SILAC channel dedicated to degradation, C_synthesis_ in the SILAC channel dedicated to synthesis. Next, Total Sum Normalization (TSN) was applied with the aim to equalize the sum of TMT peptide intensities across all TMT channels. Therefore, TMT peptide intensities from both SILAC channels were summed up to calculate one correction factor for each TMT channel:

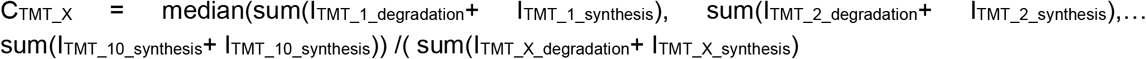

Where C_TMT_X_ is the correction factor for all peptides in the TMT channel X. TMT peptide intensities from each SILAC channel could now be corrected with the same correction factor for each TMT channel, so that their combined sum was identical between time points. Considering all protein in a TMT channel – disregarding their identiy – this fulfills the prerequisite that protein amounts stay constant over all time points.

Peptide Sum Normalization (PSN) transforms the TMT intensities of each peptide, so that the combined peptide intensity from both SILAC channels is the same across TMT channels. It thereby extends the prerequisite that protein amounts stay constant over time to the peptide level. Therefore, for each peptide a set of correction factors needs to be calculated, which normalize the intensity of every channel towards one reference channel. Which channel is designated as the reference channel is in principle arbitrary. However, we chose the 32 hour time point for this purpose because the combined SILAC light intensities were very similar to the combined SILAC heavy intensities at this time point. For each peptide the PSN correction factor can then be calculated according to:

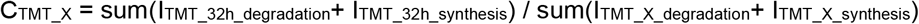

C_TMT_X_ is the correction factor for one peptide in the TMT channel X and TMT_32h denotes to the TMT channel that carries the 32 hour time point. The resulting table contained for each peptide ten correction factors, which in the case of the 32 hour timepoint was always ‘1’ (because all values had been normalized towards this time point). TMT peptide intensities from each SILAC channel could now be corrected with one set of correction factors per peptide, so that the combined sum of their TMT peptide intensities was identical between time points.

For better comparison we implemented PSN on top of TSN. Note that for the control data (‘TSN’) we present in Fig. S1 as well as Fig. S5 PSN correction factors were simply substituted by the entries ‘1’. Thereby, the control data could be treated identically to the PSN-corrected data in all of the following steps of the analysis.

In order to combine peptide synthesis curves into protein synthesis curves all TMT peptide intensities were normalized to time point infinity. Subsequently, the background at timepoint 0 was subtracted from all other timepoints. *Vice versa*, in order to combine peptide decay curves into protein decay curves all TMT peptide intensities were normalized to time point 0. Subsequently, the background at timepoint infinity was subtracted from all other timepoints. Peptides with resulting ratios smaller 0 or larger 1 at any time point other than 0 or infinity, respectively, were discarded. Normalized peptide ratios were collapsed into protein ratios in order to fit degradation or synthesis functions to them. For each timepoint the normalized peptide ratios from all peptides of a protein (‘Leading razor protein’) were collapsed into one ratio using their median. The resulting table contained the uniprot identifier of a protein and 8 (in the case of the total proteome) or 10 (in the case of the polysome profiling data) normalized protein ratios. In the synthesis case, entries for timepoint 0 were always 0 and entries for timepoint infinity always 1. In the degradation case, entries for timepoint 0 were always 1 and entries for timepoint infinity always 0. All PSN-normalized protein decay curves are reported in Table S2.

For curve fitting we used the model introduced by Boisvert et al. ^44^ for pulsed-SILAC data, which was later adapted by Welle et al. ^23^ and Zecha et al. ^24^ for TMT-SILAC. For curve fitting we used the library minpack.lm (Elzhov et al., https://CRAN.R-project.org/package=minpack.lm), which overcomes the zero-residual problem of exceptionally well-fitting data. Synthesis was modelled using the function: y ∼ (Bsyn - Asyn)*exp(-Ksyn * x) + Asyn, with the starting values Asyn = 1, Bsyn=0.3, Ksyn=0.01, the lower modelling constraints Asyn = 0, Bsyn= −2, Ksyn= −1 and the upper modelling constraints Asyn = 3, Bsyn= 2, Ksyn= 5. Degradation was modelled using the function: y ∼ (Adeg - Bdeg)*exp(-Kdeg * x) + Bdeg with the starting values Adeg = 1, Bdeg = 0.3, Kdeg = 0.01, the lower modelling constraints Adeg = 0, Bdeg= −2, Kdeg= −1 and the upper modelling constraints Adeg = 3, Bdeg= 2, Kdeg= 5. Pseudo R^2^ for the fitted functions was assessed with the library rcompanion (Mangiafico et al., https://CRAN.R-project.org/package=rcompanion), using the Nagelkerke method for comparing a non-linear model towards a null model. The null model was created by fitting a null function with the minpack.lm library for the same time points as for the actual experiment: y ∼ function(x, m){m} with the starting value m=1. Fitting results were discarded if the fitting parameters did not meet the constraints 0.5 ≤ A ≤ 2, −1 ≤ B ≤ 1, K > 0 and R^2^ ≥ 0.8 for either synthesis or degradation. Synthesis and degradation half-lives were calculated from the parameter K using the formula: HL = log(2) / K.

### Benchmarking Peptide Sum Normalization (PSN)

We benchmarked PSN using a published TMT-SILAC dataset from HeLa cells ^24^, and applied our fitting pipeline to the reported curves (supplementary table ‘Evidence_fit parameter’). Although the reported TMT PSM intensities were preprocessed we reprocessed them using our TSN and PSN pipeline. As mentioned above, this included MS3 intensity correction, total sum correction and finally peptide sum correction. For both TSN and PSN all steps were performed, yet, for TSN all peptide sum correction factors were substituted by ‘1’. Again, PSN visibly improved data continuity between time points in comparison to TSN, led to a larger number of protein half-lives for which both synthesis and decay half-lives could be determined (Figure S5B&C). Moreover, PSN improved reproducibility between the quadruplicates from an average Pearson R^2^ of 0.68 under TSN to 0.86 under PSN (Figure S5C). As mentioned in the main text, the TSN-corrected protein half-lives were similar to PSN-corrected half-lives for both synthesis and degradation, while PSN-corrected protein synthesis and decay half-lives were virtually identical (Figure S1G). We computed the relative differences between half-lives under PSN or TSN (Figure S5D) and found them in median identical for synthesis (0.99 fold) and slightly longer for decay (1.11 fold). These combined metrics confirmed the robust performance of PSN and showed that a minor distortion of half-lives occurred for some proteins in comparison to earlier approaches. In addition, we used our own protein half-life data for the MCF7 total proteome to show the effect of PSN for the individual time points in the TMT-SILAC experiment. Therefore, we deliberately applied the wrong PSN correction factors to one of the time points in our analysis. Figure S5E compares the synthesis and decay half-lives resulting from each of these analyses, demonstrating that especially erroneous correction of later time points had a stronger effect.

### Processing and Analysis of all other Proteomic Data

For the analysis of proteomic data other than TMT-SILAC data, the MaxQuant proteinGroups.txt table was used. Tables were filtered to remove ‘Potential contaminants’, ‘Reverse’ matches to the decoy database, as well as proteins ‘Only identified by site’. In all analysis, except the quantification of ribophagy upon translation inhibition, normalized protein ratios were used for SILAC quantification. For the quantification of ribophagy upon translation inhibition unnormalized protein ratios were used, which were further corrected using the four untreated replicates. Therefore, the mean of the heavy/light ratio for each protein from the untreated quadruplicates was calculated as set of correction factors. Heavy/light ratios of all proteins from the treated quadruplicates were divided by these factors individually. The average of the resulting ratios was derived and proteins filtered for a variance smaller 20 % between replicates.

For dose-response analysis of arsenite treatment on MCF7 or HeLa cells, respectively, the R package ‘drc’ ^62^ was applied.

### Functional Annotation of Proteins

For all cross-references with protein databases uniprot identifier were used. Proteins were identified as members of the cytosolic ribosome (thereby excluding mitochondrial ribosomal proteins) using the GO terms ‘cytosolic small ribosomal subunit’ and ‘cytosolic large ribosomal subunit’ and as members of the spliceosome using the GO term ‘spliceosomal complex’ in ENSEMBL BioMart (Ensembl release 96) accessed via biomaRt ^63,64^. Proteins were identified as members of particular complexes by intersection with the CORUM complex collection.

### Cryo-EM sample preparation and data collection

MCF7 cells were treated in 15 cm dishes (confluency ∼70%), by adding arsenite (Santa Cruz sc-301816) directly to the cell medium to a final concentration of 400 μM and incubating for 10 minutes at 37 °C, 5 % CO2. Lysis and polysome fractionation were performed as described above (1 dish of cells was used for each sample and loaded on one gradient), except that all fractions corresponding to 40S, 60S and 80S ribosome complexes were collected and pooled. Ribosomes were next pelleted by ultracentrifugation of the pooled fractions (which contained about 15% sucrose) for 1.5 hours at 55 000 rpm, 4 °C on a Sorvall WX90 ultracentrifuge (Beckman) with SW55Ti rotor. The supernatant was discarded, pellets were soaked in polysome buffer (50 mM Tris pH 8, 10 mM MgCl2, 140 mM KCl) for 20 minutes on ice and resuspended by pipetting. Solutions were adjusted to OD_600_≈10 and applied to carbon-coated holey copper grids (Quantifoil R2/1). Grids were glow-discharged for 30 s in oxygen/argon atmosphere of a plasma cleaner (Gatan Solarus 950) before deposition of a 4-µl aliquot of the sample. Excess buffer was blotted away for 1 s and grids were immediately vitrified in liquid ethane (−182°C) using a FEI Vitrobot (Mark II). Samples were stored in liquid nitrogen until data collection. One grid per sample was selected for data collection. Cryo-EM imaging was performed on a FEI Titan Krios 300 keV electron microscope equipped with a K3 detector (Gatan), and operated in counting mode. For the experiment with ribosomes from arsenite-naïve cells (hereinafter referred to as ‘untreated ribosomes’), illumination conditions were adjusted to collect 6,564 movie stacks, each containing 20 frames, at 1.07 Å per pixel. Illumination conditions for ribosomes from arsenite-treated cells (hereinafter referred to as ‘arsenite-treated ribosomes’) were adjusted to collect 7,112 micrographs, each containing 20 frames, at 1.07 Å per pixel. The defocus value ranged from −0.5 to 2.5 µm and the total electron dose was 27.56 e-/ Å^2^. Collection of data was automated with the EPU software (Thermo Fisher Scientific).

### Image processing

Unless stated otherwise, computational analysis of both datasets was performed using the Relion 3.1 software package ^65^. Datasets were processed separately, but following the same scheme shown in Fig. S3. The stacks of frames were aligned in MotionCor2 to correct for specimen motion during exposure, with the number of patches set to 5 × 5 ^66^. The contrast transfer function (CTF) was determined on the motion corrected sum of frames using Gctf ^67^. To generate reference templates for auto-picking, ∼1,700 particle images of ribosomes were randomly picked from the micrographs and subjected to a reference-free two-dimensional (2D) classification. No 40S subunit class average was observed among resulting 2D classes. Only 2D class averages resembling either the large 60S subunit or the full 80S ribosome were thus used as references for picking particles from the whole datasets. For validation of the auto-picking efficiency, picking of the particles was assessed manually. After this efficiency control step, the dataset with untreated ribosomes contained 1,497,356 particle images, whereas dataset with arsenite-treated ribosomes contained 1,760,231 particle images. The resulting particle images were extracted at pixel size of 4.28 Å in boxes of 128 × 128 pixels. To separate either dataset into homogenous populations, a hierarchical three-dimensional (3D) classification scheme was carried out. The initial tier of particle sorting was performed to classify the particles on the basis of global compositional heterogeneity (e.g. false positives *vs*. ribosomal species or full 80S ribosomes *vs*. 60S ribosomal subunits), before sorting based on large-scale conformational (e.g. intersubunit rotation and 40S subunit ratcheting) and subtler compositional differences (e.g. presence of ribosome inactivation factor).

For initial classification of arsenite-treated ribosomes, a human 80S ribosome (EMD-3883) ^68^ low-pass filtered to 40 Å resolution was used as a reference. One of the resulting 3D class averages (47,401 particles) showed high-resolution structural features of the 80S ribosome and was subjected to 3D auto-refinement. The refinement yielded a consensus 80S map with subnanometer global resolution, which served as internal reference for repeated initial particle sorting in either dataset. This allowed us to separate 80S ribosomes and 60S ribosomal subunits in homogenous particle groups. Only 3D class averages depicting high-resolution ribosomal features were retained for further processing, while classes containing false positives and suboptimal particles were excluded.

Well-defined features of 80S ribosomes were observed in 84,841 particles in the untreated dataset. Similarly, we identified 235,820 particles to recapitulate high-resolution features of clean 80S ribosomes in arsenite-treated dataset. All selected particles were subjected to an initial round of 3D auto-refinement using an 80S mask and 60S reference low-pass filtered to 60 Å. For this purpose, we generated a binary 80S mask using one of the best resolved 80S class averages. To generate a 60S reference, we manually segmented one of the resultant 80S class averages into 60S and 40S ribosomal subunits, and resampled the resulting 60S segment on original 80S map in UCSF Chimera ^69^. A 3D reconstruction of all untreated 80S ribosomes thus was calculated to 8.56 Å after post-processing, while the 80S reconstruction from all arsenite-treated 80S ribosomes was determined to a resolution of 8.56 Å. Following 3D auto-refinement, a single round of focused 3D classification with 40S mask and without alignment was performed. To that end, binary 40S mask was generated following the same protocol as described for 60S mask creation. This strategy was very effective at further separating clean 80S ribosomes. In total, 56,228 80S particles from untreated dataset and 53,821 80S particles from arsenite-treated dataset were retained after 3D classification. The sorted particles were re-centered, extracted at pixel size of 1.52 Å in boxes of 360 × 360 pixels and subjected to another round of 3D auto-refinement using solvent-flattened Fourier shell correlation (FSC) and otherwise standard parameters. This revealed the 80S ribosomes at 4.05 Å global resolution in untreated dataset and at 4.38 Å global resolution in arsenite-treated dataset after post-processing. Following 3D auto-refinement, particles were subjected to per-particle CTF refinement (including beam tilt estimation for individual datasets) and migrated to Relion 3.0_beta for all subsequent processing steps. The overall resolution was improved to 3.28 Å and 3.6 Å with post-processing in untreated and arsenite-treated 80S ribosomes, respectively, after Bayesian particle polishing ^70^.

To isolate homogenous particle groups within the set of 80S assemblies, we used one round of focused classification centered on the 40S subunit. The sorted particles were inspected manually and suboptimal ribosomes with a low level of interpretability were discarded. The individual classes were refined as before. To assign the best-resolved reconstructions to a particular ratcheting and rotational state of the 40S ribosomal subunit, we aligned the resultant 80S reconstructions with previously published rotated and non-rotated ribosomal structures ^29^. On the basis of particle number and map quality, in particular the 40S region, we identified two distinct states among untreated and arsenite-treated ribosomes, with the post-translocation (POST)-like state being the most abundant one in both datasets. The POST class in the untreated dataset contained 21,463 particles (38.16 %, percentage of particles from the full 80S dataset) and refined to an overall resolution of 3.77 Å after post-processing, whereas the POST class in arsenite-treated dataset was defined with 48,725 particles (90.55 %) that could be refined to an estimated resolution of 3.58 Å after post-processing. Although the 3D auto-refinement yielded cryo-EM maps at high global resolution, the small subunit density still appeared more fragmented as compared to the large subunit, suggesting further heterogeneity in this region. In order to compensate for observed local flexibility originating from intersubunit rotation and 40S subunit movement, a 3D multibody refinement was performed as described previously ^71,72^. In short, we split the 80S density into two independent segments, comprising the 40S subunit and the 60S subunit. This strategy resulted in 40S density segments at 3.72 Å (untreated ribosomes) and 3.65 Å (arsenite-treated ribosomes) global resolution after post-processing using the 40S mask also employed for 3D multibody refinement. Similarly, the results yielded 60S density segments at 3.09 Å (untreated ribosomes) and 3.28 Å (arsenite-treated ribosomes) after post-processing with the 60S mask also used for 3D multibody refinement.

To calculate compositional and conformational differences between the POST-like 80S ribosome in either dataset and a reference human ribosome in POST state (EMD-10674), we applied the difference map approach using TEMPy ^73^. The maps were first low-pass filtered to 6 Å. The difference between maps was calculated after map-to-map fitting and based on global scaling. The locations and a shape of the beak region within the 40S subunit and the P-stalk area within the 60S subunit were identified as the difference peaks in both datasets (Fig. 2K and Fig. S6K).

In parallel, 80S assemblies were subjected to independent rounds of focused 3D classification with a mask focusing on factors that were visible at low threshold levels already in the consensus refinement structure, i.e. either LYAR or IFRD2, to enrich for their occupancy. This enabled us to identify one LYAR-enriched class with 7,314 particles in the untreated dataset and one LYAR-containing class with 8,242 particles from the arsenite-treated dataset. Each resultant class was refined using the 3D auto-refinement procedure against the respective particles within that class with a soft reference mask in the shape of the 80S ribosome, thus yielding reconstructions at 5.64 Å and 6.08 Å global resolution after post-processing in the untreated and arsenite-treated dataset, respectively. Similarly, a total of 9,487 80S untreated 80S ribosomes and 6,985 arsenite-treated 80S ribosomes were found to depict the strongest density of IFRD2, and were determined to a resolution of 4.34 Å (untreated IFRD2-80S complex) and 5.82 Å (arsenite-treated IFRD2-80S complex) after post-processing.

Finally, for processing of the 60S subunits, we used a similar protocol as described above. All processing steps preceding Bayesian particle polishing were performed in Relion 3.1. Briefly, after initial 3D classification, well-defined features of 60S subunits were observed in 43,436 particles among untreated ribosomal assemblies. In parallel, we identified 324,519 particles to show high-resolution features of clean 60S subunits among the arsenite-treated ribosomal assemblies. In an attempt to improve resolution of the 60S subunit, we extracted pre-selected 60S particles at full spatial resolution (1.07 Å, box size 384 × 384 pixels) and subjected them to 3D auto-refinement. After 3D auto-refinement, particles were subjected to CTF refinement transferred to Relion 3.0_beta for Bayesian particle polishing using a training set of 5,000 particles. The resulting ‘shiny’ particles were subjected to a final round of 3D auto-refinement. For post-processing, a solvent mask was generated from the final map from 3D auto-refinement, low-pass filtered to 15 Å. The initial binary mask was extended by 5 Å in all directions and a raised-cosine edge was added to create a soft mask. A final refinement of 60S particles resulted in post-processed 3.14 Å (untreated 60S) and 2.85 Å (arsenite-treated 60S) cryo-EM maps. Additionally, focused 3D classification was performed on the eIF6 binding area of the 60S ribosomal subunits to enrich for its occupancy. Starting with 43,436 untreated particles, multiple rounds of focused 3D classification were used to isolate two classes of 13,004 particles in which the eIF6 density was best resolved. The two classes were pooled and refined together to 3.84 Å after post-processing, with a mask encompassing the full eIF6-60S complex.

The global resolution and the temperature factors of the maps were estimated by applying a soft mask around the ribosome density and using the “gold standard” FSC criterion of independently refined half maps (FSC = 0.143) within Relion.

### Gene-Ontology (GO) Enrichment Analysis

Ranked GO enrichment analysis was performed using the GOrilla ^74^ web interface and uniprot identifier as input.

### Statistical Analysis and Data Visualization

Data was filtered by variance cut-offs, which were individually selected for each experiment with the objective to include as many data points as possible, while excluding outliers or highly variable data that would undermine statistical power. All data handling apart from what is mentioned above was performed in R (3.5.3) with RStudio (1.1.463) and visualized using the ggplot2 ^75^ library. Figures were arranged in Illustrator (CC, Adobe 2015).

## Notes

### Competing Interest Statement

The authors have declared no competing interest.

## References

1. Dikic, I.Proteasomal and Autophagic Degradation Systems. Annu. Rev. Biochem. 86, 193–224 (2017).

2. Wiśniewski, J. R., Hein, M. Y., Cox, J. & Mann, M. A ‘proteomic ruler’ for protein copy number and concentration estimation without spike-in standards. Mol. Cell. Proteomics 3497–3506 (2014). doi:10.1074/mcp.M113.037309

3. Sung, M.-K., Reitsma, J. M., Sweredoski, M. J., Hess, S. & Deshaies, R. J. Ribosomal proteins produced in excess are degraded by the ubiquitin–proteasome system. Mol. Biol. Cell 27, 2642–2652 (2016).

4. Sung, M. K. et al. A conserved quality-control pathway that mediates degradation of unassembled ribosomal proteins. Elife 5, 1–28 (2016).

5. Lam, Y. W., Lamond, A. I., Mann, M. & Andersen, J. S. Analysis of Nucleolar Protein Dynamics Reveals the Nuclear Degradation of Ribosomal Proteins. Curr. Biol. 17, 749–760 (2007).

6. Yanagitani, K., Juszkiewicz, S. & Hegde, R. S. UBE2O is a quality control factor for orphans of multiprotein complexes. Science (80-.). 357, 472–475 (2017).

7. Peña, C., Hurt, E. & Panse, V. G. Eukaryotic ribosome assembly, transport and quality control. Nat. Struct. Mol. Biol. 24, 689–699 (2017).

8. Jackson, R. J., Hellen, C. U. T. & Pestova, T. V. The mechanism of eukaryotic translation initiation and principles of its regulation. Nat. Rev. Mol. Cell Biol. 11, 113–127 (2010).

9. Dever, T. E. & Green, R. The Elongation, Termination, and Recycling Phases of Translation in Eukaryotes. Cold Spring Harb. Perspect. Biol. 4, a013706–a013706 (2012).

10. Pisarev, A. V., Hellen, C. U. T. & Pestova, T. V. Recycling of eukaryotic posttermination ribosomal complexes. Cell 131, 286–99 (2007).

11. Pisarev, A. V. et al. The Role of ABCE1 in Eukaryotic Posttermination Ribosomal Recycling. Mol. Cell 37, 196–210 (2010).

12. Khatter, H. et al. Purification, characterization and crystallization of the human 80S ribosome. Nucleic Acids Res. 42, e49–e49 (2014).

13. Khatter, H., Myasnikov, A. G., Natchiar, S. K. & Klaholz, B. P. Structure of the human 80S ribosome. Nature 520, 640–645 (2015).

14. Kraft, C., Deplazes, A., Sohrmann, M. & Peter, M. Mature ribosomes are selectively degraded upon starvation by an autophagy pathway requiring the Ubp3p/Bre5p ubiquitin protease. Nat. Cell Biol. 10, 602–610 (2008).

15. Gretzmeier, C. et al. Degradation of protein translation machinery by amino acid starvation-induced macroautophagy. Autophagy 13, 1064–1075 (2017).

16. Kristensen, A. R. et al. Ordered Organelle Degradation during Starvation-induced Autophagy. Mol. Cell. Proteomics 7, 2419–2428 (2008).

17. An, H., Ordureau, A., Körner, M., Paulo, J. A. & Harper, J. W. Systematic quantitative analysis of ribosome inventory during nutrient stress. Nature 583, 303–309 (2020).

18. Wyant, G. A. et al. Nufip1 is a ribosome receptor for starvation-induced ribophagy. Science (80-.). 360, 751–758 (2018).

19. Liu, Y. et al. Autophagy-dependent rRNA degradation is essential for maintaining nucleotide homeostasis during C. elegans development. Elife 7, e36588 (2018).

20. Trendel, J. et al. The Human RNA-Binding Proteome and Its Dynamics during Translational Arrest. Cell 176, 391–403.e19 (2019).

21. Imami, K. et al. Phosphorylation of the Ribosomal Protein RPL12/uL11 Affects Translation during Mitosis. Mol. Cell 1–15 (2018). doi:10.1016/j.molcel.2018.08.019

22. Yap, M. G. S. et al. Multitagging Proteomic Strategy to Estimate Protein Turnover Rates in Dynamic Systems. J. Proteome Res. 9, 2087–2097 (2010).

23. Welle, K. A. et al. Time-resolved Analysis of Proteome Dynamics by Tandem Mass Tags and Stable Isotope Labeling in Cell Culture (TMT-SILAC) Hyperplexing. Mol. Cell. Proteomics 15, 3551–3563 (2016).

24. Zecha, J. et al. Peptide Level Turnover Measurements Enable the Study of Proteoform Dynamics. Mol. Cell. Proteomics 17, 974–992 (2018).

25. Zylber, E. A. & Penman, S. The effect of high ionic strength on monomers, polyribosomes, and puromycin-treated polyribosomes. Biochim. Biophys. Acta - Nucleic Acids Protein Synth. 204, 221–229 (1970).

26. Martin, T. E. & Hartwell, L. H. Resistance of active yeast ribosomes to dissociation by KCl. J. Biol. Chem. 245, 1504–6 (1970).

27. Thoms, M. et al. Structural basis for translational shutdown and immune evasion by the Nsp1 protein of SARS-CoV-2. Science (80-.). 369, 1249–1256 (2020).

28. Brown, A., Baird, M. R., Yip, M. C., Murray, J. & Shao, S. Structures of translationally inactive mammalian ribosomes. Elife 7, 1–18 (2018).

29. Behrmann, E. et al. Structural snapshots of actively translating human ribosomes. Cell 161, 845–857 (2015).

30. Shao, S. et al. Decoding Mammalian Ribosome-mRNA States by Translational GTPase Complexes. Cell 167, 1229–1240.e15 (2016).

31. Zhou, Y. et al. Structural impact of K63 ubiquitin on yeast translocating ribosomes under oxidative stress. Proc. Natl. Acad. Sci. U. S. A. 117, 22157–22166 (2020).

32. Kedersha, N. L., Gupta, M., Li, W., Miller, I. & Anderson, P. RNA-Binding Proteins Tia-1 and Tiar Link the Phosphorylation of Eif-2α to the Assembly of Mammalian Stress Granules. J. Cell Biol. 147, 1431–1442 (1999).

33. Huang, M.-T. Harringtonine, an Inhibitor of Initiation of Protein Biosynthesis. Mol. Pharmacol. 11, 511–519 (1975).

34. Fresno, M., Jimenez, A., Vazquez, D., Jiménez, A. & Vázquez, D. Inhibition of Translation in Eukaryotic Systems by Harringtonine. Eur. J. Biochem. 72, 323–330 (1977).

35. Ingolia, N. T., Lareau, L. F. & Weissman, J. S. Ribosome profiling of mouse embryonic stem cells reveals the complexity and dynamics of mammalian proteomes. Cell 147, 789–802 (2011).

36. Nathans, D. PUROMYCIN INHIBITION OF PROTEIN SYNTHESIS: INCORPORATION OF PUROMYCIN INTO PEPTIDE CHAINS. Proc. Natl. Acad. Sci. 51, 585–592 (1964).

37. Gao, X. et al. Quantitative profiling of initiating ribosomes in vivo. Nat. Methods 12, 147–153 (2015).

38. Liu, B. & Qian, S.-B. Characterizing inactive ribosomes in translational profiling. Translation 4, e1138018 (2016).

39. Eichelbaum, K., Winter, M., Diaz, M. B., Herzig, S. & Krijgsveld, J. Selective enrichment of newly synthesized proteins for quantitative secretome analysis. Nat. Biotechnol. 30, 984–990 (2012).

40. Eichelbaum, K. & Krijgsveld, J. Rapid temporal dynamics of transcription, protein synthesis, and secretion during macrophage activation. Mol. Cell. Proteomics 13, 792–810 (2014).

41. An, H. & Harper, J. W. Systematic analysis of ribophagy in human cells reveals bystander flux during selective autophagy. Nat. Cell Biol. 20, 135–143 (2018).

42. Kong, A. T., Leprevost, F. V, Avtonomov, D. M., Mellacheruvu, D. & Nesvizhskii, A. I. MSFragger: ultrafast and comprehensive peptide identification in mass spectrometry–based proteomics. Nat. Publ. Gr. 293, (2017).

43. Budkevich, T. V. et al. Regulation of the mammalian elongation cycle by subunit rolling: A eukaryotic-specific ribosome rearrangement. Cell 158, 121–131 (2014).

44. Boisvert, F.-M. et al. A Quantitative Spatial Proteomics Analysis of Proteome Turnover in Human Cells. Mol. Cell. Proteomics 11, M111.011429 (2012).

45. McShane, E. et al. Kinetic Analysis of Protein Stability Reveals Age-Dependent Degradation. Cell 167, 803–815.e21 (2016).

46. Mancera-Martínez, E., Brito Querido, J., Valasek, L. S., Simonetti, A. & Hashem, Y. ABCE1: A special factor that orchestrates translation at the crossroad between recycling and initiation. RNA Biol. 14, 1279–1285 (2017).

47. Pisareva, V. P., Skabkin, M. A., Hellen, C. U. T., Pestova, T. V. & Pisarev, A. V. Dissociation by Pelota, Hbs1 and ABCE1 of mammalian vacant 80S ribosomes and stalled elongation complexes. EMBO J. 30, 1804–1817 (2011).

48. Van Den Elzen, A. M. G., Schuller, A., Green, R. & Séraphin, B. Dom34-Hbs1 mediated dissociation of inactive 80S ribosomes promotes restart of translation after stress. EMBO J. 33, 265–276 (2014).

49. Murakami, R. et al. The Interaction between the Ribosomal Stalk Proteins and Translation Initiation Factor 5B Promotes Translation Initiation. Mol. Cell. Biol. 38, (2018).

50. Delarue, M. et al. mTORC1 Controls Phase Separation and the Biophysical Properties of the Cytoplasm by Tuning Crowding. Cell 174, 338–349.e20 (2018).

51. Ceci, M. et al. Release of eIF6 (p27BBP) from the 60S subunit allows 80S ribosome assembly. Nature 426, 579–584 (2003).

52. Gribling-Burrer, A. S. et al. A dual role of the ribosome-bound chaperones RAC/Ssb in maintaining the fidelity of translation termination. Nucleic Acids Res. 47, 7018–7034 (2019).

53. Thoreen, C. C. et al. A unifying model for mTORC1-mediated regulation of mRNA translation. Nature 485, 109–113 (2012).

54. Simonetti, A., Guca, E., Bochler, A., Kuhn, L. & Hashem, Y. Structural Insights into the Mammalian Late-Stage Initiation Complexes. Cell Rep. 31, 107497 (2020).

55. Voorhees, R. M. & Hegde, R. S. Structures of the scanning and engaged states of the mammalian srp-ribosome complex. Elife 4, 1–21 (2015).

56. Schmidt, C. et al. The cryo-EM structure of a ribosome–Ski2-Ski3-Ski8 helicase complex. Science (80-.). 354, 1431–1433 (2016).

57. Bhaskar, V. et al. Dynamics of uS19 C-Terminal Tail during the Translation Elongation Cycle in Human Ribosomes. Cell Rep. 31, 107473 (2020).

58. Brown, A., Shao, S., Murray, J., Hegde, R. S. & Ramakrishnan, V. Structural basis for stop codon recognition in eukaryotes. Nature 524, 493–496 (2015).

59. Hughes, C. S. et al. Ultrasensitive proteome analysis using paramagnetic bead technology. Mol. Syst. Biol. 10, 757 (2014).

60. Bertolini, M. et al. Interactions between nascent proteins translated by adjacent ribosomes drive homomer assembly. Science (80-.). 371, (2021).

61. Cox, J. & Mann, M. MaxQuant enables high peptide identification rates, individualized p.p.b.-range mass accuracies and proteome-wide protein quantification. Nat. Biotechnol. 26, 1367–72 (2008).

62. Ritz, C., Baty, F., Streibig, J. C. & Gerhard, D. Dose-response analysis using R. PLoS One 10, 1–13 (2015).

63. Zerbino, D. R. et al. Ensembl 2018. Nucleic Acids Res. 46, D754–D761 (2018).

64. Durinck, S., Spellman, P. T., Birney, E. & Huber, W. Mapping identifiers for the integration of genomic datasets with the R/ Bioconductor package biomaRt. Nat. Protoc. 4, 1184–1191 (2009).

65. Zivanov, J. et al. New tools for automated high-resolution cryo-EM structure determination in RELION-3. Elife 7, 1–22 (2018).

66. Zheng, S. Q. et al. MotionCor2: Anisotropic correction of beam-induced motion for improved cryo-electron microscopy. Nat. Methods 14, 331–332 (2017).

67. Zhang, K. Gctf: Real-time CTF determination and correction. J. Struct. Biol. 193, 1–12 (2016).

68. Natchiar, S. K., Myasnikov, A. G., Kratzat, H., Hazemann, I. & Klaholz, B. P. Visualization of chemical modifications in the human 80S ribosome structure. Nature 551, 472–477 (2017).

69. Pettersen, E. F. et al. UCSF Chimera - A visualization system for exploratory research and analysis. J. Comput. Chem. 25, 1605–1612 (2004).

70. Zivanov, J., Nakane, T. & Scheres, S. H. W. A Bayesian approach to beam-induced motion correction in cryo-EM single-particle analysis. IUCrJ 6, 5–17 (2019).

71. Wild, K. et al. MetAP-like Ebp1 occupies the human ribosomal tunnel exit and recruits flexible rRNA expansion segments. Nat. Commun. 11, 1–10 (2020).

72. Nakane, T., Kimanius, D., Lindahl, E. & Scheres, S. H. W. Characterisation of molecular motions in cryo-EM single-particle data by multi-body refinement in RELION. Elife 7, 1–18 (2018).

73. Farabella, I. et al. TEMPy: A Python library for assessment of three-dimensional electron microscopy density fits. J. Appl. Crystallogr. 48, 1314–1323 (2015).

74. Eden, E., Navon, R., Steinfeld, I., Lipson, D. & Yakhini, Z. GOrilla: a tool for discovery and visualization of enriched GO terms in ranked gene lists. BMC Bioinformatics 10, 48 (2009).

75. Wickham, H. ggplot2. (Springer International Publishing, 2016). doi:10.1007/978-3-319-24277-4

76. Klinge, S., Voigts-Hoffmann, F., Leibundgut, M., Arpagaus, S. & Ban, N. Crystal structure of the eukaryotic 60S ribosomal subunit in complex with initiation factor 6. Science (80-.). 334, 941–948 (2011).

