## Supplemental figures for "Translational Activity Controls Ribophagic Flux and Turnover of Distinct Ribosome Pools"

**Supplementary Materials:**

Figs. S1: Proteomic quantification of MCF7 polysome profiling fraction and comparison of normalization procedures for TMT-SILAC data.

Figs. S2: Benchmarking peptide sum normalization.

Figs. S3: Cryo-EM processing workflow for ribosomal complexes with and without arsenite treatment.

Figs. S4: Local resolution estimation and Fourier shell correlation curves.

Figs. S5: Identification of LYAR and IFRD2 bound to translationally inactive 80S ribosomes without and with arsenite treatment.

Figs. S6: Cryo-EM structure of the inactive human 80S ribosome after arsenite treatment.

Figs. S7: Arsenite treatment constrains conformational plasticity of the inactive 80S ribosome.

Figs. S8: Identification of eIF6 bound to 60S ribosomal subunits without arsenite treatment.

Table S1: Label-free quantification of proteins in MCF7 polysome profiling fractions.

Table S2: Protein half-lives from the MCF7 total proteome and MCF7 polysome profiling fractions.

Table S3: Changes in the MCF7 total proteome upon translation inhibition.

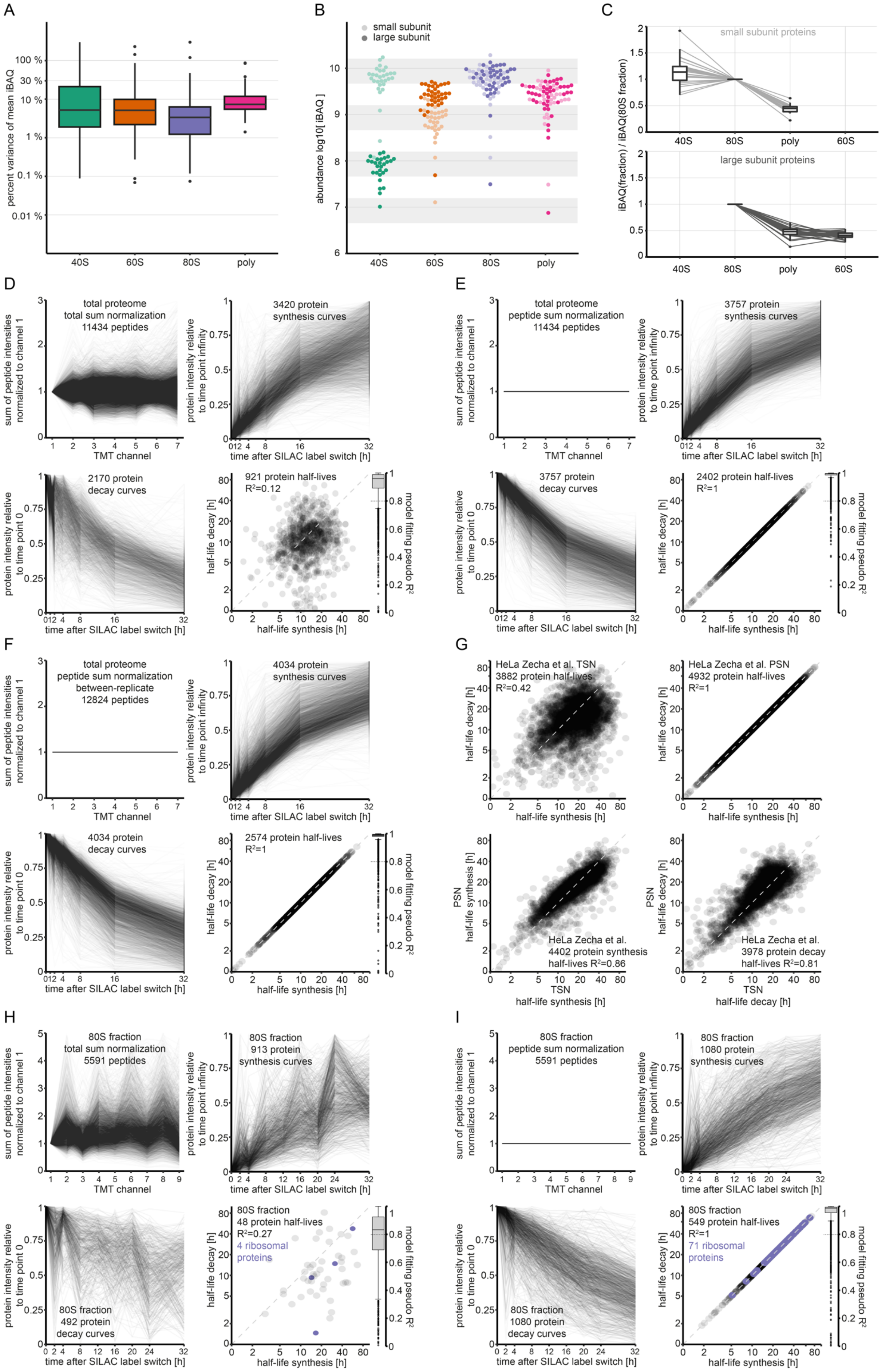
A
percent variance of mean iBAQ
100 %
30 %
10 %
1 %
0.1 %
0.01 %
40S
60S
80S
poly
B
small subunit
large subunit
abundance log10[ iBAQ ]
C
small subunit proteins
large subunit proteins
iBAQ(fraction) / iBAQ(80S fraction)
D
total proteome
total sum normalization
11434 peptides
sum of peptide intensities normalized to channel 1
TMT channel
3420 protein synthesis curves
protein intensity relative to time point infinity
time after SILAC label switch [h]
2170 protein decay curves
protein intensity relative to time point 0
921 protein half-lives
R²=0.12
half-life decay [h]
half-life synthesis [h]
model fitting pseudo R²
E
total proteome
peptide sum normalization
11434 peptides
3757 protein synthesis curves
3757 protein decay curves
2402 protein half-lives
R²=1
F
total proteome
peptide sum normalization
between-replicate
12824 peptides
4034 protein synthesis curves
4034 protein decay curves
2574 protein half-lives
R²=1
G
HeLa Zecha et al. TSN
3882 protein half-lives
R²=0.42
HeLa Zecha et al. PSN
4932 protein half-lives
R²=1
PSN half-life synthesis [h]
HeLa Zecha et al.
4402 protein synthesis half-lives R²=0.86
TSN half-life synthesis [h]
PSN half-life decay [h]
HeLa Zecha et al.
3978 protein decay half-lives R²=0.81
TSN half-life decay [h]
H
80S fraction
total sum normalization
5591 peptides
80S fraction
913 protein synthesis curves
80S fraction
492 protein decay curves
80S fraction
48 protein half-lives
R²=0.27
4 ribosomal proteins
I
80S fraction
peptide sum normalization
5591 peptides
80S fraction
1080 protein synthesis curves
80S fraction
1080 protein decay curves
80S fraction
549 protein half-lives
R²=1
71 ribosomal proteins

**Fig. S1: Proteomic quantification of MCF7 polysome profiling fraction and comparison of normalization procedures for TMT-SILAC data.** A) Boxplot comparing variances of cytosolic ribosomal protein abundances within polysome profiling fractions from MCF7 cells. Triplicates of MCF7 cells were fractionated as indicated in Figure 1B. B) Dotplot showing absolute contribution of ribosomal proteins within each polysome profiling fraction. Light shading refers to small, dark shading to large subunit proteins. Displayed are proteins with a variance in iBAQ between biological triplicates smaller 30 %. Grey shading illustrates the maximal standard deviation around powers of ten assuming a variance of 30 %. C) Assembly plot comparing the relative stoichiometry of ribosomal proteins between polysome profiling fractions of MCF7 cells. Compared are iBAQ values of cytosolic ribosomal proteins normalized to the 80S fraction. Ratios were calculated from the mean iBAQ of triplicate experiments for proteins with a variance smaller 30 %. Boxplots visualize the distribution of relative abundances within each fraction, and lines connect individual proteins between the fraction. Top: Small subunit proteins assembling into 80S or polysome fractions. Bottom: Large subunit proteins assembling into 80S or polysome fractions. For visibility large subunit proteins are omitted from the 40S fraction and small subunit proteins from the 60S fraction. D-F) Analysis of TMT-SILAC data from the total proteome of MCF7 cells for the computation of protein half-lives using different normalization methods. Combined data from both biological replicates are shown. Compared are total sum normalization (TSN, D), peptide sum normalization (PSN, E) and peptide sum normalization using the other biological replicate (F). Top left panel: Lineplot displaying the sum of TMT peptide intensities from both SILAC channels (SILAC light + SILAC heavy) for all peptides in the analysis. For comparison intensities are normalized to time point 0. Top right panel: Lineplot showing protein synthesis curves. Bottom left panel: Lineplot showing protein decay curves. Bottom right panel: Scatterplot comparing protein synthesis and decay half-lives. $R^2$ refers to the Pearson correlation between synthesis and decay. Boxplot on the right side shows the distribution of pseudo $R^2$ values achieved during the model fitting of all protein synthesis and decay curves to the model. For details see Materials and Methods. G) Analysis of published TMT-SILAC data from the total proteome of HeLa cells [24] for the computation of protein half-lives using different normalization methods. Data from four biological replicates is shown. Protein synthesis ot decay half-lives, respectively, that could be derived from more than one replicate were averaged. Top left panel: Scatterplot comparing protein synthesis and decay half-lives derived from total sum normalized data (TSN). Top right panel: Scatterplot comparing protein synthesis and decay half-lives derived from peptide sum normalized data (PSN). Bottom left panel: Scatterplot comparing protein synthesis half-lives between TSN and PSN normalized data. Bottom right panel: Scatterplot comparing protein decay half-lives between TSN and PSN normalized data. $R^2$ refers to the Pearson correlation. For further details see Figures S5A-D and Materials and Methods. H&I) Analysis of TMT-SILAC data from polysome profiling of MCF7 cells for the computation of protein half-lives using different normalization methods. Exemplary data for the 80S fraction is presented. Same panel display as in D-F.

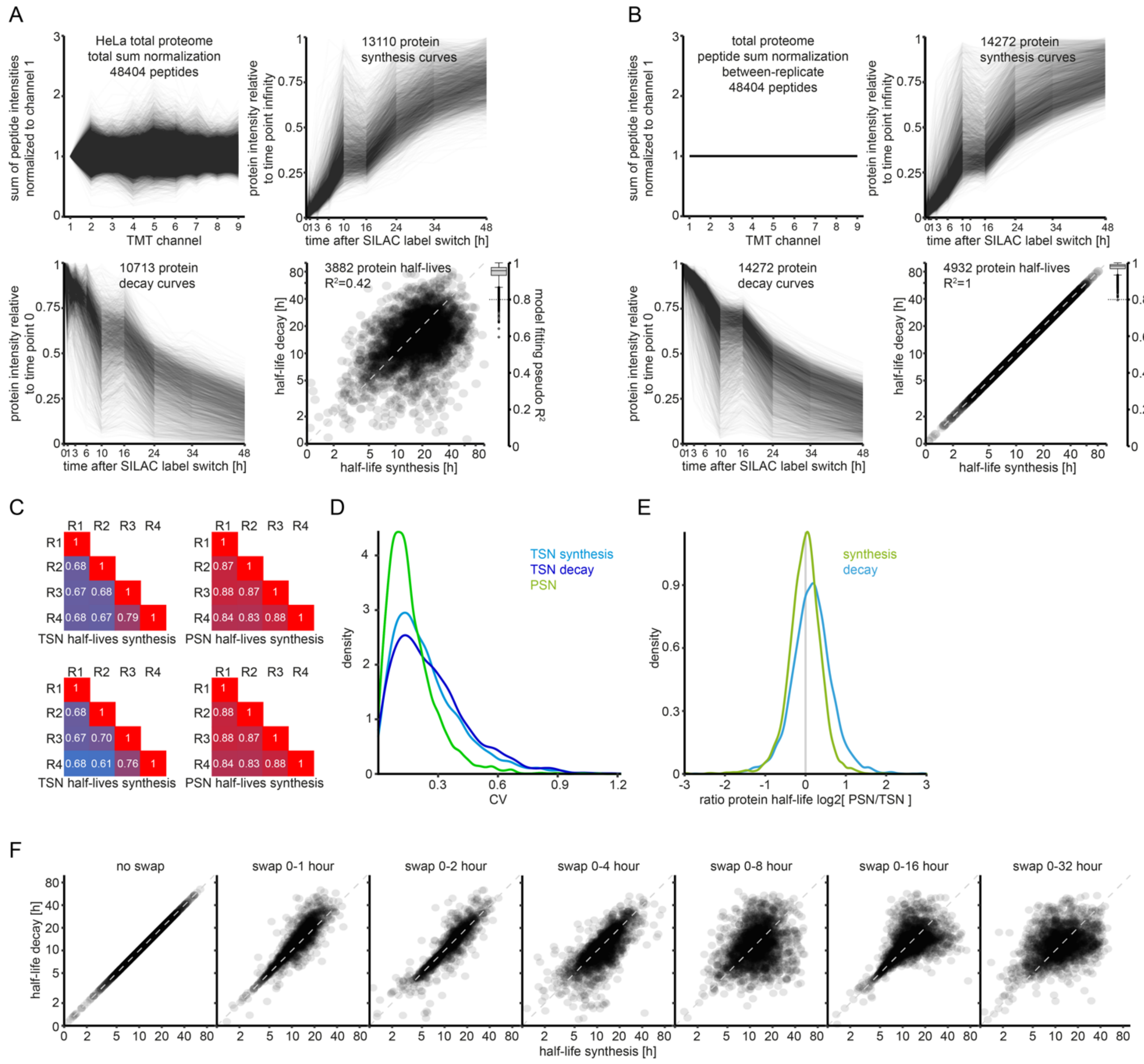


**Fig. S2: Benchmarking peptide sum normalization.** A-D) TMT-SILAC data from HeLa cells reported by Zecha et al. was reanalyzed using our analysis pipeline for the computation of protein half-lives. See Materials and Methods for details. A) Analysis of published synthesis and decay curves for the computation of protein half-lives using different normalization methods. Combined data from all four biological replicates are shown. Compared are total sum normalization (TSN, A) and peptide sum normalization (PSN, B). Top left panel: Lineplot displaying the sum of TMT peptide intensities from both SILAC channels (SILAC light + SILAC heavy) for all peptides in the analysis. For comparison intensities are normalized to time point 0. Top right panel: Lineplot showing protein synthesis curves. Bottom left panel: Lineplot showing protein decay curves. Bottom right panel: Scatterplot comparing protein synthesis and decay half-lives. $R^2$ refers to the Pearson correlation between synthesis and decay. Boxplot on the right side shows the distribution of pseudo $R^2$ values achieved during the model fitting of all protein synthesis and decay curves to the model. For details see Materials and Methods. C) Heatmaps showing the Pearson correlation of synthesis (top) or decay (bottom) half-lives between biological replicates, when TSN (left) or PSN (right) was applied. D) Density plot comparing the coefficient of variation (CV) of half-lives derived from quadruplicates with PSN or TSN. CVs for synthesis and decay are identical under PSN. E) Density plot comparing PSN to TSN-derived half-lives. See also Figure S1G. F) Control analysis of TMT-SILAC data from the total proteome of MCF7 cells for the computation of protein half-lives using swapped PSN correction factors. Analysis occurred as in Figure S1E but PSN correction factors for the indicated time point were exchanged for PSN correction factors of the (wrong) time point 0.

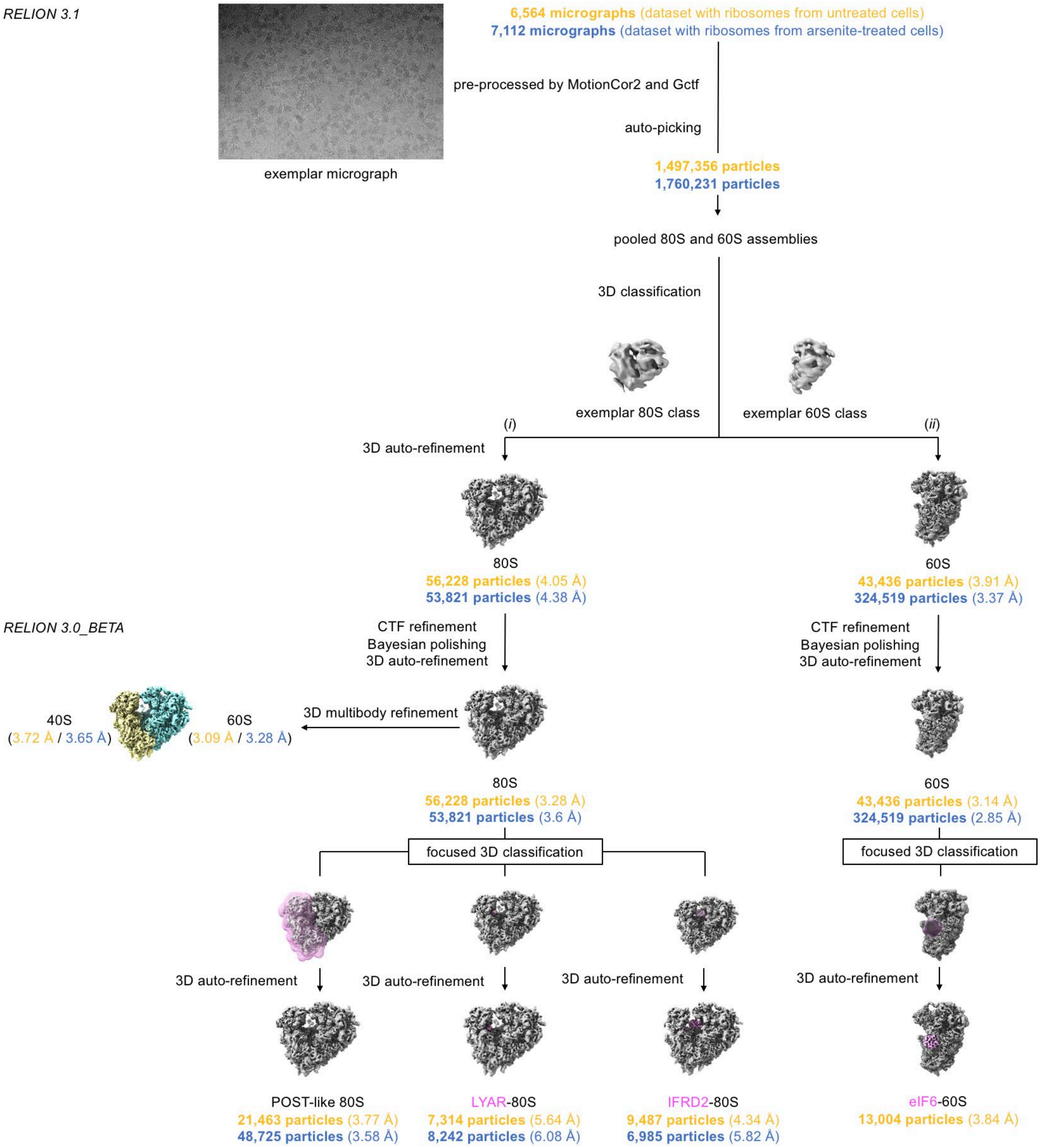


**Fig. S3: Cryo-EM processing workflow for ribosomal complexes with and without arsenite treatment.** The diagram depicts schematically the steps of computational analysis for the cryo-EM datasets of 80S ribosomes (i) and 60S ribosomal subunits (ii) from untreated (yellow) and arsenite-treated cells (blue). Software packages, programs and individual procedure steps are listed, together with micrograph and particle numbers. The final reconstructions correspond to five cryo-EM densities in each dataset: consensus 80S ribosome, POST-like 80S ribosome, LYAR-80S complex, IFRD2-80S complex, and 60S subunit. For untreated cells, the eIF6-60S complex was also analysed. Additional information is available in the Materials and Methods section.

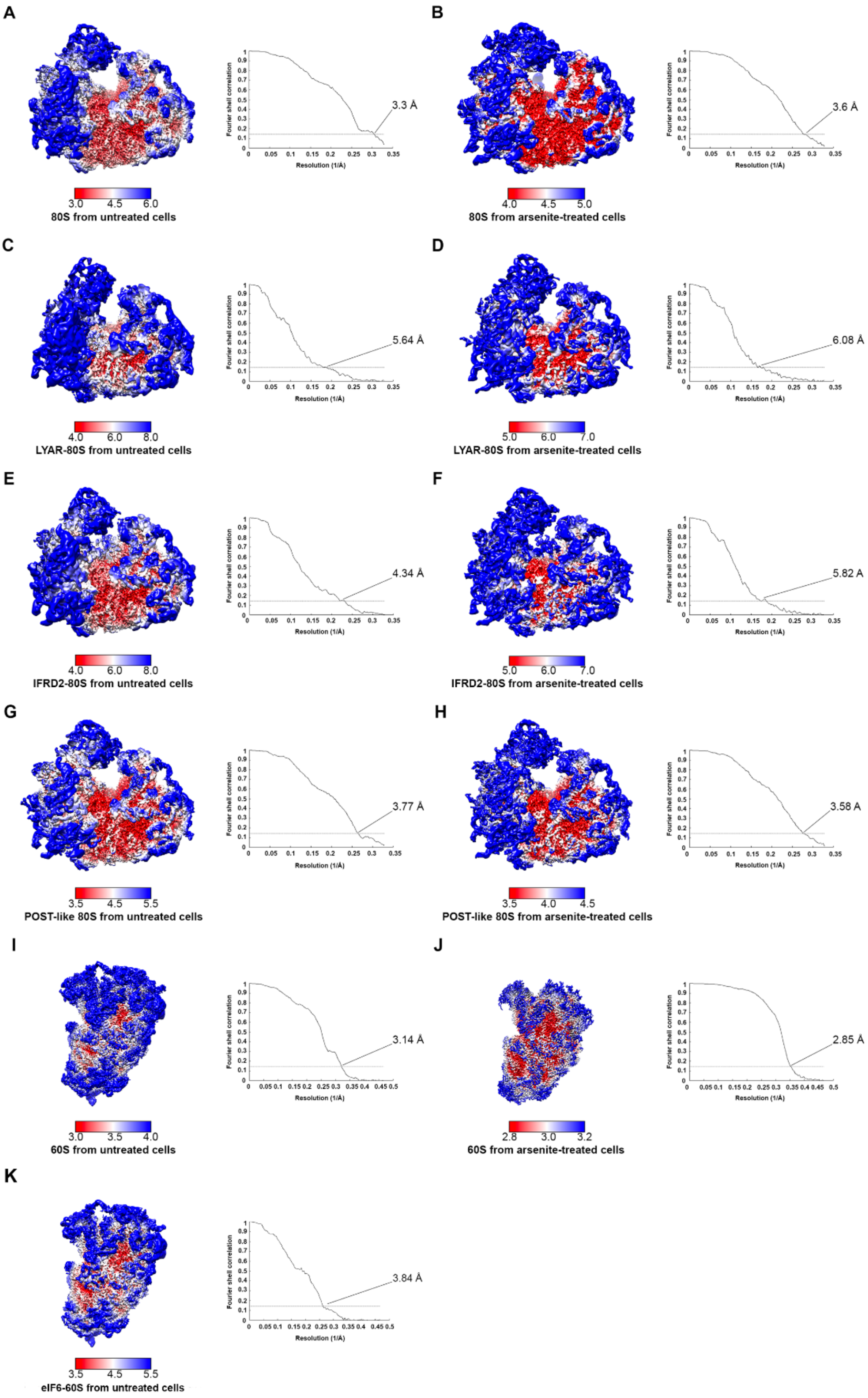
A
3.3 Å
3.0 4.5 6.0
80S from untreated cells
B
3.6 Å
4.0 4.5 5.0
80S from arsenite-treated cells
C
5.64 Å
4.0 6.0 8.0
LYAR-80S from untreated cells
D
6.08 Å
5.0 6.0 7.0
LYAR-80S from arsenite-treated cells
E
4.34 Å
4.0 6.0 8.0
IFRD2-80S from untreated cells
F
5.82 A
5.0 6.0 7.0
IFRD2-80S from arsenite-treated cells
G
3.77 Å
3.5 4.5 5.5
POST-like 80S from untreated cells
H
3.58 A
3.5 4.0 4.5
POST-like 80S from arsenite-treated cells
I
3.14 Å
3.0 3.5 4.0
60S from untreated cells
J
2.85 Å
2.8 3.0 3.2
60S from arsenite-treated cells
K
3.84 Å
3.5 4.5 5.5
eIF6-60S from untreated cells
Fourier shell correlation
Resolution (1/Å)

**Fig. S4: Local resolution estimation and Fourier shell correlation curves.** (A - K) Left: Final cryo-EM densities filtered and colored according to local resolution. Right: Fourier shell correlation (FSC) curves between two independently refined half maps. Threshold for final resolution estimation according to the 'gold-standard' set at FSC = 0.143 and indicated by a dashed line.

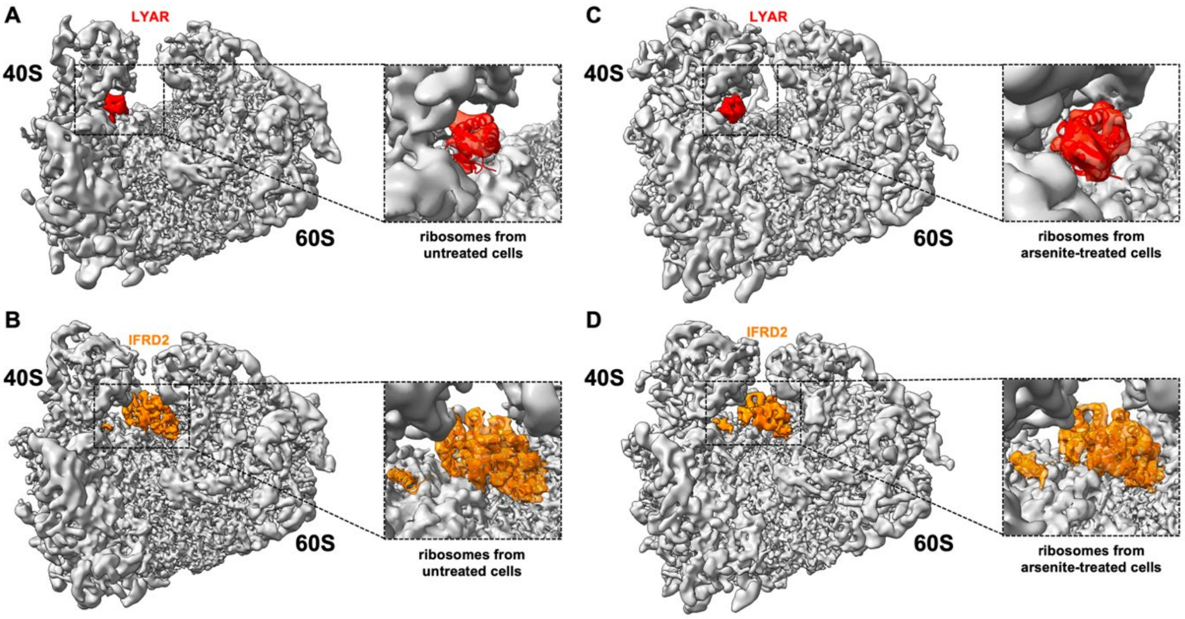


**Fig. S5: Identification of LYAR and IFRD2 bound to translationally inactive 80S ribosomes without and with arsenite treatment.** A) Cryo-EM density of the inactive human LYAR-80S complex without arsenite treatment at 5.64 Å resolution. In the zoomed view, the atomic model of LYAR was superimposed together with the structure of an inactive human 80S ribosome (PDB-6ZMO) [27]. B) Same as A, but the cryo-EM density at 6.08 Å resolution was obtained from arsenite-treated cells. C) Cryo-EM density of the inactive human IFRD2-80S complex without arsenite treatment at 4.34 Å resolution. In the zoomed view, the atomic model of IFRD2 was superimposed with the structure of an inactive human 80S ribosome (PDB-6MTC) [28]. D) Same as C, but the cryo-EM density at 5.82 Å resolution was obtained from arsenite-treated cells. The LYAR and IFRD2 densities are displayed at a different contour level than the ribosome maps.

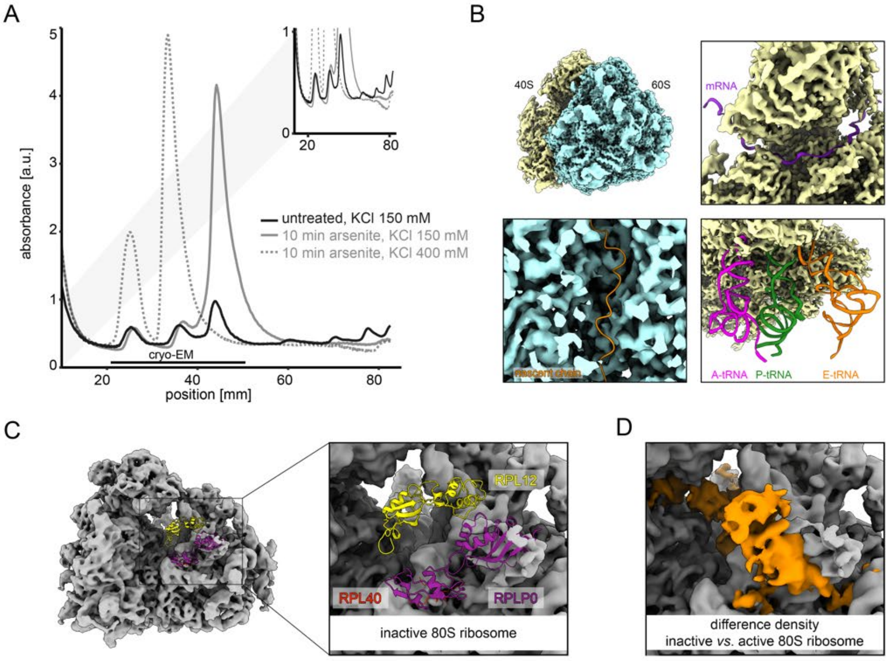


**Fig. S6: Cryo-EM structure of the inactive human 80S ribosome after arsenite treatment.** A) Polysome profiling of MCF7 cells after 10 minutes of arsenite stress (grey) under low and high salt conditions. For comparison the polysome profiling UV trace of untreated cells is overlayed (black). B) Top left: local-resolution-filtered cryo-EM density of inactive human 80S ribosomes under arsenite stress after 3D multi-body refinement. Large (blue; 3.28 Å global resolution) and small (yellow; 3.65 Å global resolution) ribosomal subunits are indicated. 60S, large 60S ribosomal subunit; 40S, small 40S ribosomal subunit. Top right: the mRNA channel is devoid of density for an mRNA molecule (PDB-6YAL)[54]. Bottom left: density of the 60S subunit cut through the polypeptide exit tunnel. The ribosomal exit tunnel site is devoid of density for a nascent polypeptide (PDB-3JAJ)[55]. E) Bottom right: the intersubunit space is vacant and devoid of density for tRNAs (PDB-5MC6 and PDB-5LZV)[30,56]. C) Cryo-EM density of inactive human ribosome lowpass filtered to 6-Å resolution for clarity. Atomic models for the components of the P-stalk area have been superposed (PDB-3JAH)[58]. (D) Zoomed view of positive difference density (in orange) calculated by the subtraction of the reconstruction shown in (C) from the reconstruction of active human ribosome lowpass filtered to 6-Å resolution shown in Fig. 2B (EMD-10674)[57]. Parts of (C) colored as previously.

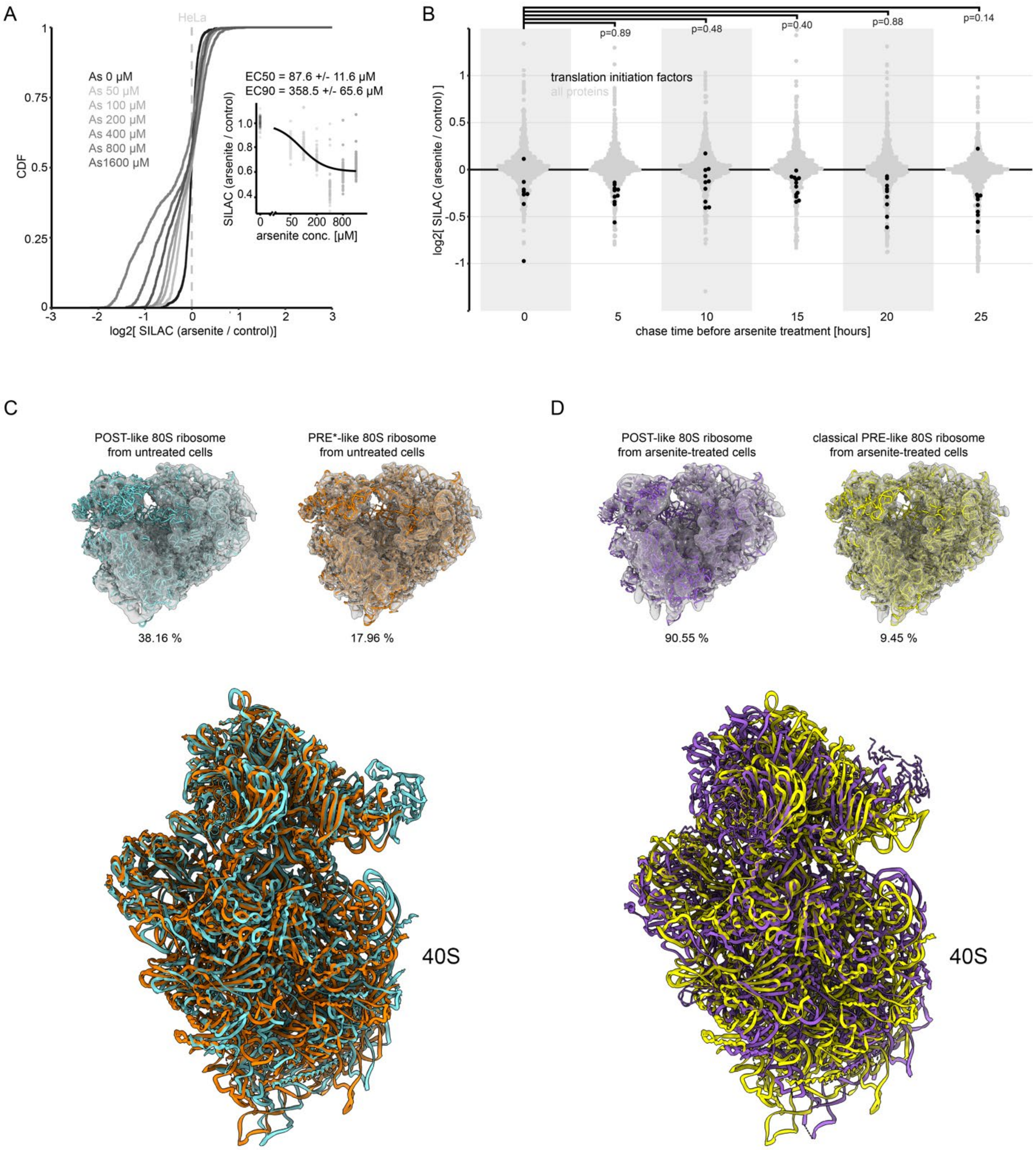


**Fig. S7: Arsenite treatment constrains conformational plasticity of the inactive 80S ribosome.** A) Cumulative distribution of expression changes in the total proteome of HeLa cells upon increasing doses of arsenite added for 30 minutes of treatment. Reproduction of experiment in Figure 4A with HeLa instead of MCF7 cells. B) Dotplot comparing arsenite-induced autophagy within MCF7 proteomes of different ages. Same as in Figure 4C but black dots represent proteins with the GO annotation 'translation initiation factor activity, RNA-binding'. Light grey dots represent all proteins in the analysis. Shown are means of duplicate experiments with label swap filtered for a variance smaller 20 %. C) Top: Cryo-EM densities of the POST-like and PRE*-like 80S ribosomes without arsenite treatment at 3.77 Å and 4.41 Å resolution, respectively. The structures were superimposed with the atomic models of the human 80S ribosome in POST-state (orange, PDB-6Y2L) or hybrid PRE-state (turquoise, PDB-6Y57)[57]. Relative abundances of the classes are given. Bottom: overlay of the two atomic models according to the 60S subunit rRNA highlights the movement of the 40S ribosomal subunit against the 60S ribosomal subunit. The superposed models are shown in a frontal view on the 40S ribosomal subunit and the overlapping 60S ribosomal subunits are not shown for clarity. D) Top: Cryo-EM densities of the POST-like and classical-1 PRE-like 80S ribosomes with arsenite treatment at 3.58 Å and 6.22 Å resolution, respectively. The structures were superimposed with the atomic models of the human 80S ribosome in POST-state (purple; PDB-6Y2L) or classical-PRE-state (yellow; PDB-6Y0G)[57]. Relative abundances of the classes are given. Bottom: same as in C.

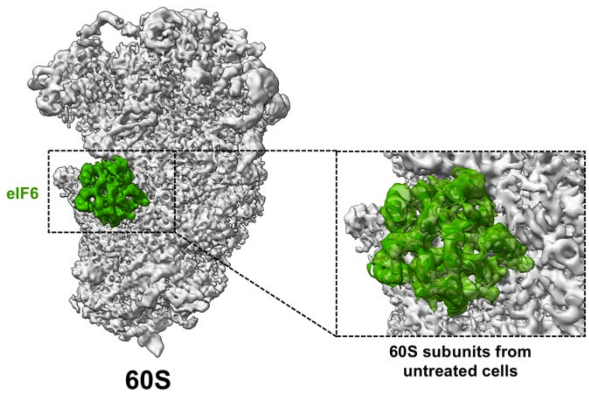


**Fig. S8: Identification of eIF6 bound to 60S ribosomal subunits without arsenite treatment.** Cryo-EM density of the human eIF6-60S complex without arsenite treatment at 3.84 Å resolution. In the zoomed view, the atomic model of eIF6 was superimposed together with the structure of a human 60S ribosomal subunit (PDB-4V8P)[76]. The eIF6 density is displayed at a different contour level than the ribosome density.
